# β-Catenin-NFκB-CFTR interactions in cholangiocytes regulate inflammation and fibrosis during ductular reaction

**DOI:** 10.1101/2021.09.15.460429

**Authors:** Shikai Hu, Jacquelyn O. Russell, Silvia Liu, Ravi Rai, Karis Kosar, Junyan Tao, Edward Hurley, Minakshi Poddar, Sucha Singh, Aaron Bell, Donghun Shin, Reben Raeman, Aatur D. Singhi, Kari Nejak-Bowen, Sungjin Ko, Satdarshan P. Monga

## Abstract

Expansion of biliary epithelial cells (BECs) during ductular reaction (DR) is observed in liver diseases including cystic fibrosis (CF), and associated with inflammation and fibrosis, *albeit* without complete understanding of underlying mechanism. Using two different genetic knockouts of β-catenin, one with β-catenin loss is hepatocytes and BECs (KO1), and another with loss in only hepatocytes (KO2), we demonstrate disparate long-term repair after an initial injury by 2-week choline-deficient ethionine-supplemented diet. KO2 show gradual liver repopulation with BEC-derived β-catenin- positive hepatocytes, and resolution of injury. KO1 showed persistent loss of β-catenin, NF-κB activation in BECs, progressive DR and fibrosis, reminiscent of CF histology. We identify interactions of β-catenin, NFκB and CF transmembranous conductance regulator (CFTR) in BECs. Loss of CFTR or β-catenin led to NF-κB activation, DR and inflammation. Thus, we report a novel β-catenin-NFκB-CFTR interactome in BECs, and its disruption may contribute to hepatic pathology of CF.

## Introduction

The liver possesses unique regenerative potential. During chronic liver injury, however, liver fibrosis accompanies regeneration and can progress to cirrhosis, which can then progress to end-stage liver disease (ESLD) or hepatocellular cancer (HCC)^1^. Currently, cirrhosis is the 11^th^ leading cause of death globally, and the incidence of liver disease continues to rise as conditions such as non-alcoholic fatty liver disease (NAFLD) and alcoholic liver disease continue to prevail^2^. Thus, there has been great interest in studying mechanisms of injury, inflammation and fibrosis during liver injury in order to effectively develop novel therapies. The role of hepatic epithelial cells (referred henceforth as ‘hepithelial’ cells), which include both hepatocytes and cholangiocytes or biliary epithelial cells (BECs), in regulating microenvironment is beginning to be appreciated. Loss of hepatocyte differentiation in chronic liver diseases and ESLD, either due to much needed hepithelial proliferation for repair, or as an adaptation to escape injury, seems to contribute to not only loss of key hepatic functions, but is also causally associated with increased immune response and hepatic fibrosis^3, 4^. However, how hepithelial cells may modulate hepatic immune microenvironment, is unclear.

As an important hepithelial cell type, BECs are known to undergo proliferation to replace dying BECs in cholangiopathies or cystic liver diseases, as well as can under phenotypic switch to generate *de novo* hepatocytes when hepatocytes are chronically injured and/or are unable to optimally proliferate, phenomena termed as ductular reaction (DR)^5, 6^. Reactive ductules, however, can secrete pro-inflammatory and pro-fibrotic cytokines to induce inflammation, activate myofibroblasts, and induce fibrosis.

The extent of DR correlates with fibrosis in many types of liver injuries^7,8,9^. The molecular underpinnings of reactive DR is incompletely understood although molecules like Yes associated protein-1 (YAP1) have been implicated^10^.

β-Catenin, the major downstream effector of the Wnt signaling is a well-known mediator of hepatocyte proliferation. Liver-specific (hepatocyte and BECs) β-catenin knockout (KO1) mice generated by breeding β-catenin-floxed and albumin-cre mice show delayed liver regeneration (LR) after partial hepatectomy or after toxicant-induced liver injury^11, 12^. When KO1 were administered choline-deficient, ethionine-supplemented (CDE) diet, it triggered greater steatosis, cell death, DR, inflammation and fibrosis than wild-type (WT1), and upon switching to normal diet for 2 weeks (2w) for recovery, continued to show greater injury due to an impairment of hepatocyte proliferation^13, 14^. Similar greater injury, fibrosis and DR was observed in CDE-fed hepatocyte-only β- catenin KO (KO2), generated by delivering adeno-associated virus serotype 8 carrying a plasmid encoding *Cre* recombinase under a hepatocyte-specific thyroxine-binding globulin (TBG) promoter (AAV8-TBG-Cre) into the β-catenin-floxed mice^14^. Intriguingly, labeling BECs for fate-tracing, showed the liver repair to occur through BEC-to-hepatocyte transdifferentiation upon recovery for 2w and up to 6 months on normal diet, although long-term impact on injury resolution, inflammation, DR and fibrosis was not studied in either model^14^.

In the current study, we investigate hepatic injury and repair in KO2 and KO1 mice challenged for 2w with CDE diet and allowed to recover on normal diet for 2w, 3 months (3m) and 6m. Intriguingly, we observed highly divergent injury-repair responses in the two models. KO2 mice showed progressive repair through expansion of BEC- derived β-catenin-positive hepatocytes, and resolution of inflammation, DR and fibrosis.

However, KO1 display progressive and peculiar DR composed of numerous small luminal structures lined by a single layer of BECs, even after being on normal diet for 6m, which is associated with fibrosis and inflammation, and is reminiscent of Cystic fibrosis (CF)-like morphology. We identify a unique interactome of β-catenin, p65 subunit of NF-κB and Cystic fibrosis transmembranous conductance regulator (CFTR) in BECs and show perturbations in these interactions leading to excessive NF-κB activation and inflammation in BECs in both KO1 and CF patients.

## Results

### Long term follow-up of mice lacking β-catenin in hepatocytes only (KO2) show delayed but eventual resolution of fibrosis and DR after initial 2w CDE diet

We previously showed CDE diet for 2w led to enhanced injury, fibrosis, and DR in mice lacking β-catenin in hepatocytes only (KO2), generated by delivering AAV8-TBG-Cre into *Ctnnb1^flox/flox^; Rosa-stop^flox/flox^-EYFP* mice, as compared to WT2 mice, generated by injecting AAV8-TBG-Cre into *Ctnnb1^+/+^; Rosa-stop^flox/flox^-EYFP* mice^14^. And that upon switching to normal diet, liver repair occurred via hepatocyte proliferation in WT2 but through BEC-to-hepatocyte transdifferentiation in KO2^14^. To specifically investigate durability of repair especially after the increased injury observed in the KO2 mice at 2w of CDE diet, we fed CDE diet to KO2 and WT2 mice for 2w and switched to normal diet for 2w, 3m or 6m (Fig.1A). KO2 mice had elevated serum alanine aminotransferase (ALT) and total bilirubin (BR) levels than WT2 at 2w of CDE diet, but returned to normal at 2w onwards after switching to normal diet, similar to WT2, although BR levels tended to be higher in KO2 up to 3m of recovery (Fig.1B). Alkaline phosphatase (ALP) was increased in WT2 and KO2 after 2w of CDE injury but returned to normal at 2w of recovery in both groups (Fig.1B).

**Figure 1:**
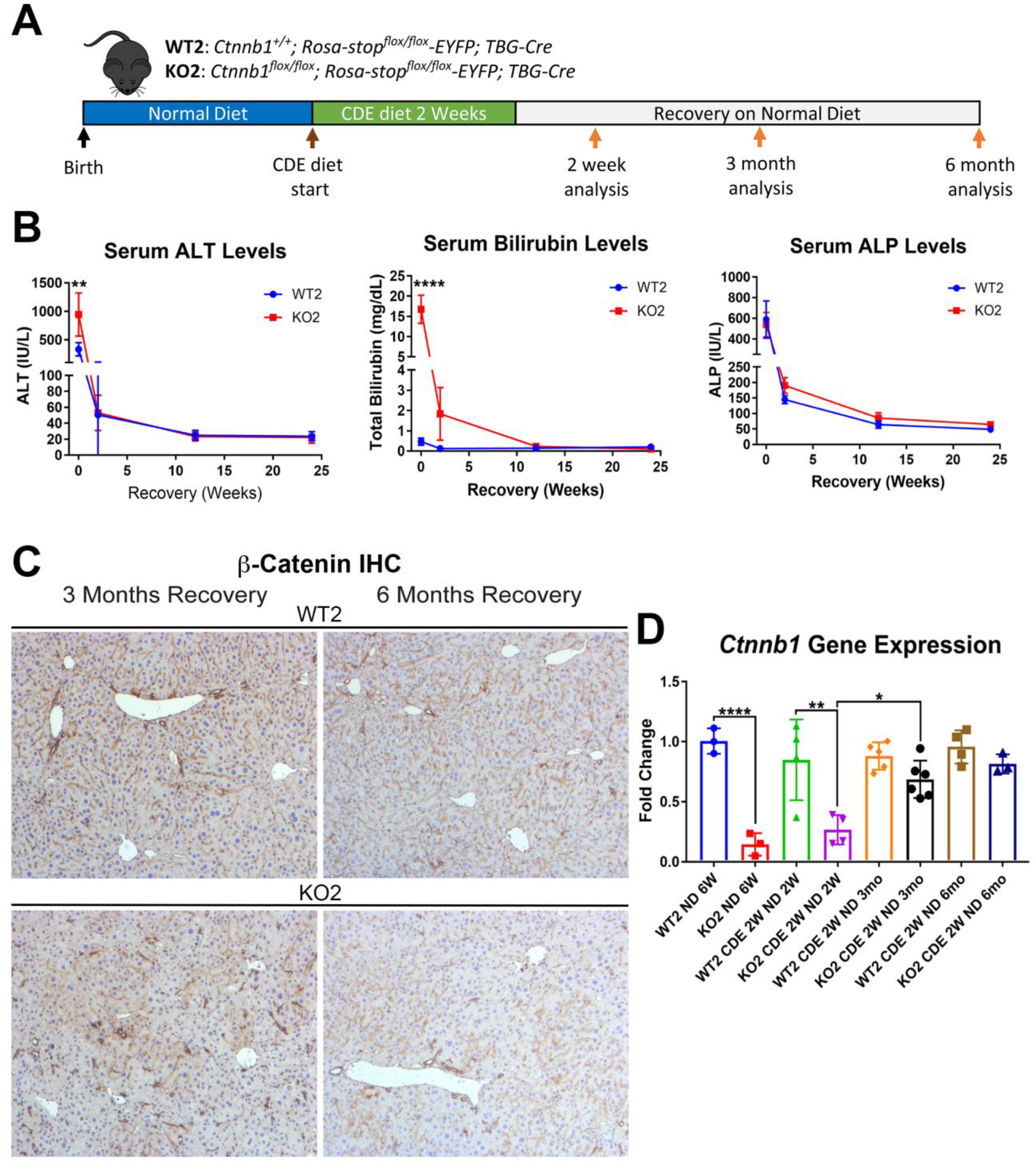
Comparable recovery of WT2 and KO2 on normal diet after initial 2w CDE diet injury, along with repopulation of KO2 livers with BEC-derived β-catenin- positive hepatocytes. A) Experimental design showing WT2 and KO2 on 2w of CDE diet and recovery on normal diet for up to 6m with analysis at intermediate time-points as indicated. B) Serum ALT, bilirubin, and alkaline phosphatase (ALP) in the two groups over time (one-way ANOVA, **p<0.01, ****p<0.0001, n = 3 to 6 per group). C) β-Catenin immunohistochemistry in WT2 and KO2 mice at 3m and 6m of recovery showing β- catenin-positive BECs and hepatocytes in KO2 and WT2. Scale bar = 50µm. D) *Ctnnb1* gene expression in WT2 and KO2 mice during recovery from CDE diet (one-way ANOVA, *p<0.05; **p<0.01, ****p<0.0001. n = 3 to 6 per group. Individual animal values represented by dots.)

Previously by fate-tracing, we observed a BECs transdifferentiated to hepatocytes and expanded in KO2^14^. Likewise, by immunohistochemistry (IHC) for β- catenin that is only present in BECs in KO2 at baseline, there were increased numbers of β-catenin-positive hepatocytes at 3m and 6m of recovery (Fig.1C). RT-PCR for β- catenin gene (*Ctnnb1)* expression showed increasing expression in KO2 livers over time, becoming comparable to WT2 at 6m (Fig.1D).

Sirius red staining for fibrosis showed greater collagen deposition in KO2 than WT2 at 2w of CDE diet and persisted at 2w of recovery (Fig.2A,2B). Interestingly, despite being on normal diet and lack of any ongoing injury, KO2 continued to show fibrosis at 3m, eventually resolving at 6m (Fig. 2A, 2B). Likewise, expression of *Col1a1* and *Tgfβ2* tended to be higher in KO2 compared to WT2 mice at 3m recovery, but were comparable to WT2 at 6m (Fig.2C).

**Figure 2:**
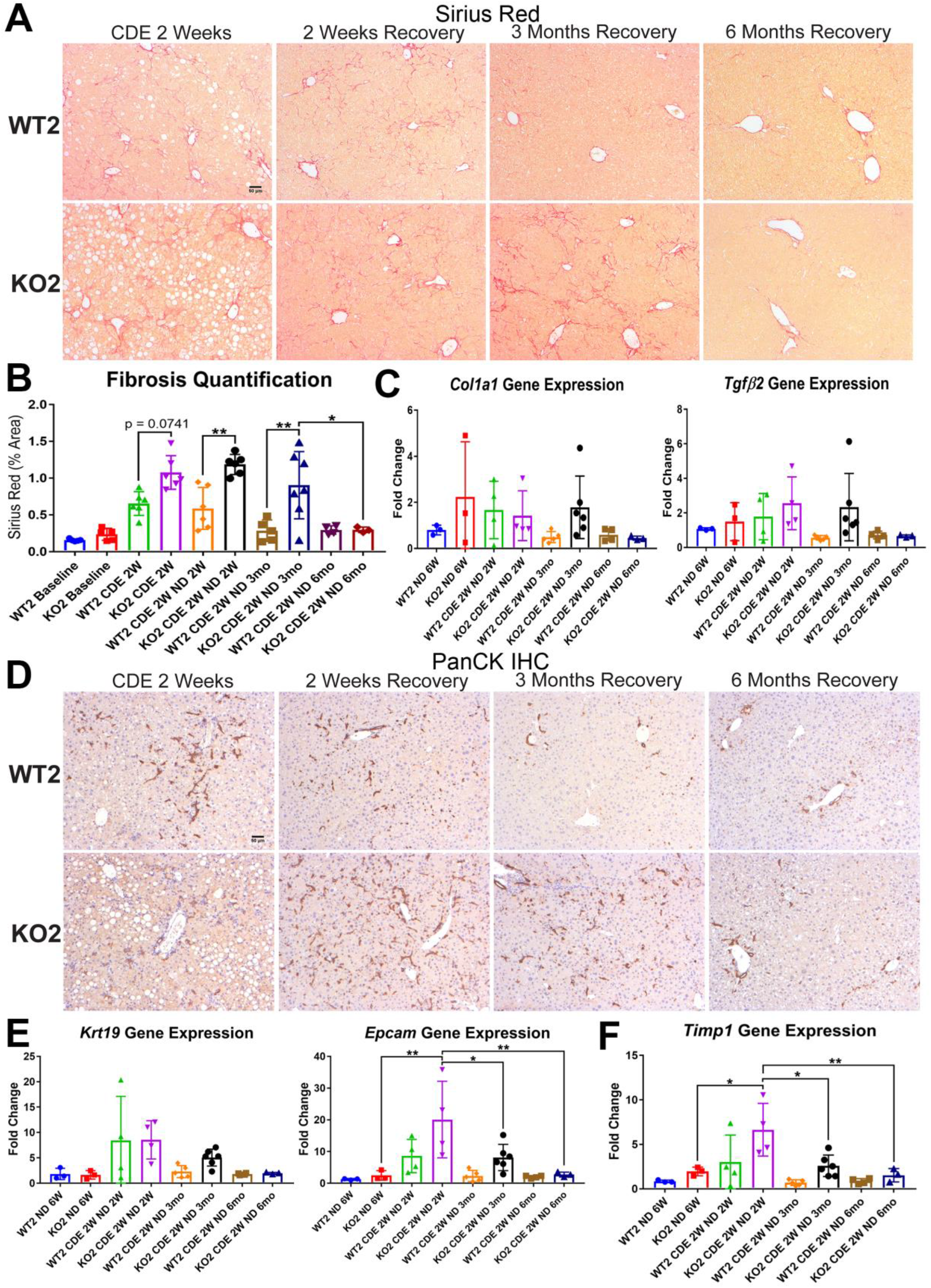
Fibrosis and DR is sustained in KO2 mice recovering on normal diet until 3m, but subsides by 6m, after initial 2w CDE diet. A) Sirius Red staining in WT2 and KO2 mice over time during recovery from CDE diet. Scale bar = 50µm. B) Quantification of Sirius Red staining (one way-ANOVA, *p<0.05, **p<0.01. n = 3 to 7 per group. Individual animal values represented by dots.). C) A trend of increased expression of *Col1a1* and *Tgfβ2* in KO2 mice at 3m of recovery but not at 6m (n = 3 to 6 per group. Individual animal values represented by dots.). D) Pan-cytokeratin (PanCK) staining in WT2 and KO2 mice over time during recovery from CDE diet. Scale bar = 50 µm. E) A trend of higher *Krt19* expression and significantly higher expression of *Epcam* gene in KO2 up to 3m on recovery and normalization to WT2 levels at 6m (one-way ANOVA, *p<0.05, **p<0.01. n = 3 to 6 per group. Individual animal values represented by dots.). F) Significantly higher *Timp1* gene expression in KO2 than WT2 up to 3m on recovery diet and normalization at 6m (one-way ANOVA, *p<0.05, **p<0.01. n = 3 to 6 per group. Individual animal values represented by dots.)

Since increased DR was observed in KO2 after CDE-diet induced injury, and fibrosis can be associated with DR, we next performed IHC for pan-cytokeratin (PanCK) (Fig.2D). There was robust DR in KO2 mice and WT2 mice after 2w CDE diet, and was also evident at 2w of recovery although it was more pronounced in KO2. At 3m of recovery, normal bile ducts are seen in WT2 whereas DR composed of flattened, non-luminal, and single or few cell clusters is evident throughout liver lobule in KO2 (Fig.2D). At 6m, there was no DR in either group (Fig.2D). Gene expression of BEC markers *Krt19* and *Epcam* confirmed these observations (Fig.2E).

Expression of the gene encoding tissue inhibitor of metalloproteinase 1a (*Timp1*), a well-known inhibitor of matrix metalloproteinases, known for a role in extracellular matrix degradation, was determined next as a possible mechanism of fibrosis resolution^15, 16^. Higher expression of *Timp1* persisted in KO2 as compared to WT2 at all recovery times except 6m, coinciding with resolution of fibrosis and DR (Fig.2F).

Taken together, these results suggest that higher DR is associated with greater fibrosis in KO2, and resolution of the DR and fibrosis took longer in KO2 than WT2, which correlated with enhanced repopulation of the KO2 liver with β-catenin-positive hepatocytes and normalization of Timp1 levels.

### Long term follow-up of mice lacking β-catenin in hepatocytes and BECs (KO2) show prolonged fibrosis and DR without any evidence of regression after initial 2w CDE-diet

Next, we next placed *Albumin-Cre^+/-^ Ctnnb1^flox/flox^* (KO1) mice lacking β- catenin in hepatocytes and BECs, and their wild-type littermates (WT1) on CDE diet for 2w and allowed recovery on normal diet for 2w, 3m and 6m (Fig.3A). We observed severe liver injury in KO1 mice, shown by significantly higher serum ALT and total BR after 2w of CDE diet compared to WT1. During recovery, serum ALT levels in KO1 and WT1 mice decreased to normal levels (Fig.3B), while BR remained mildly elevated in KO1 mice up to 3m of recovery as compared to WT1 which returned to normal at 2w of recovery (Fig.3B). Serum ALP levels were comparably increased in WT1 and KO1 at 2w of CDE injury and returned to normal levels at 2w recovery (Fig.3B).

**Figure 3:**
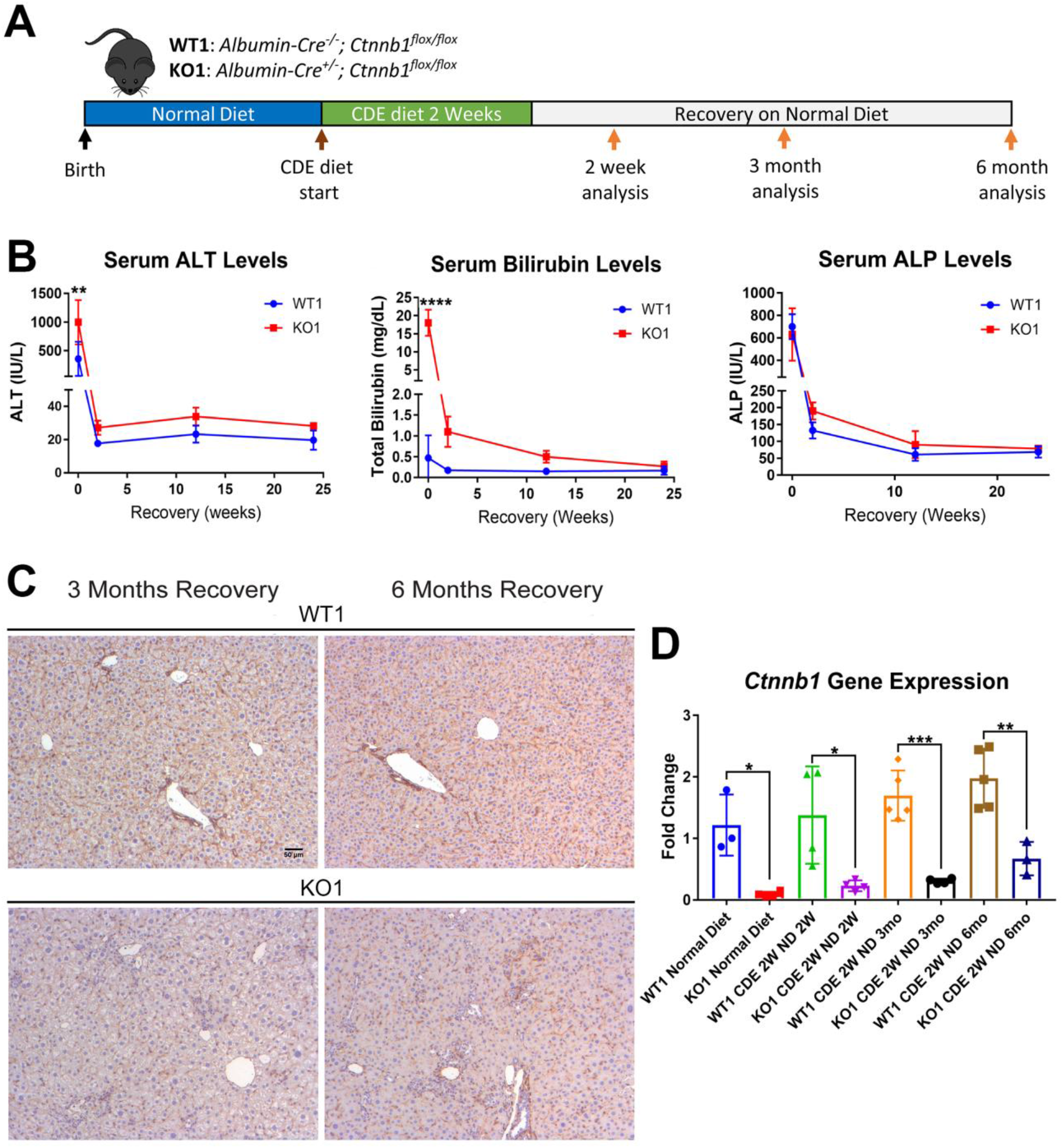
Serum biochemistry suggests comparable recovery on normal diet in WT1 and KO1 after 2w CDE diet and continued lack of β-catenin in KO1. A) Experimental design showing WT1 and KO1 on 2 weeks of CDE diet and recovery on normal diet for up to 6m with analysis at intermediate time-points as indicated. B) Serum ALT, bilirubin, and ALP in the two groups over time. (One-way ANOVA, **p<0.01, ****p<0.0001, n = 3 to 5 per group). C) β-Catenin immunohistochemistry in WT1 and KO1 mice at 3m and 6m of recovery showing absence of β-catenin in BECs and hepatocytes in KO1. Scale bar = 50µm. D) *Ctnnb1* gene expression in WT1 and KO1 mice during recovery from CDE diet shows continued β-catenin absence over time in KO1 (one-way ANOVA, *p<0.05; **p<0.01, ****p<0.0001. n = 3 to 5 per group. Individual animal values represented by dots.)

Since β-catenin is lacking in hepithelial cells in the KO1 livers, IHC for β-catenin and RT-PCR for *Ctnnb1* showed continued absence in KO1 and not WT1 at 3m and 6m recovery on normal diet (Fig.3C,3D).

We previously reported increased fibrosis in KO1 mice compared to WT1 littermates after 2w CDE diet^14^. Here, we evaluated fibrosis during recovery on normal diet in both WT1 and KO1. Despite normalization of serum transaminases during recovery, we observed continued fibrosis especially in the periportal area in KO1 especially at 3m and 6m by Sirius Red staining, whereas WT1 mice displayed resolution of fibrosis as early as 2w recovery (Fig.4A). Quantification verified significant increases in fibrosis in KO1 at all time points compared to WT1 (Fig.4B). Additionally, expression of *Col1a1* tended to be higher in KO1 mice during recovery (Fig.4C).

**Figure 4:**
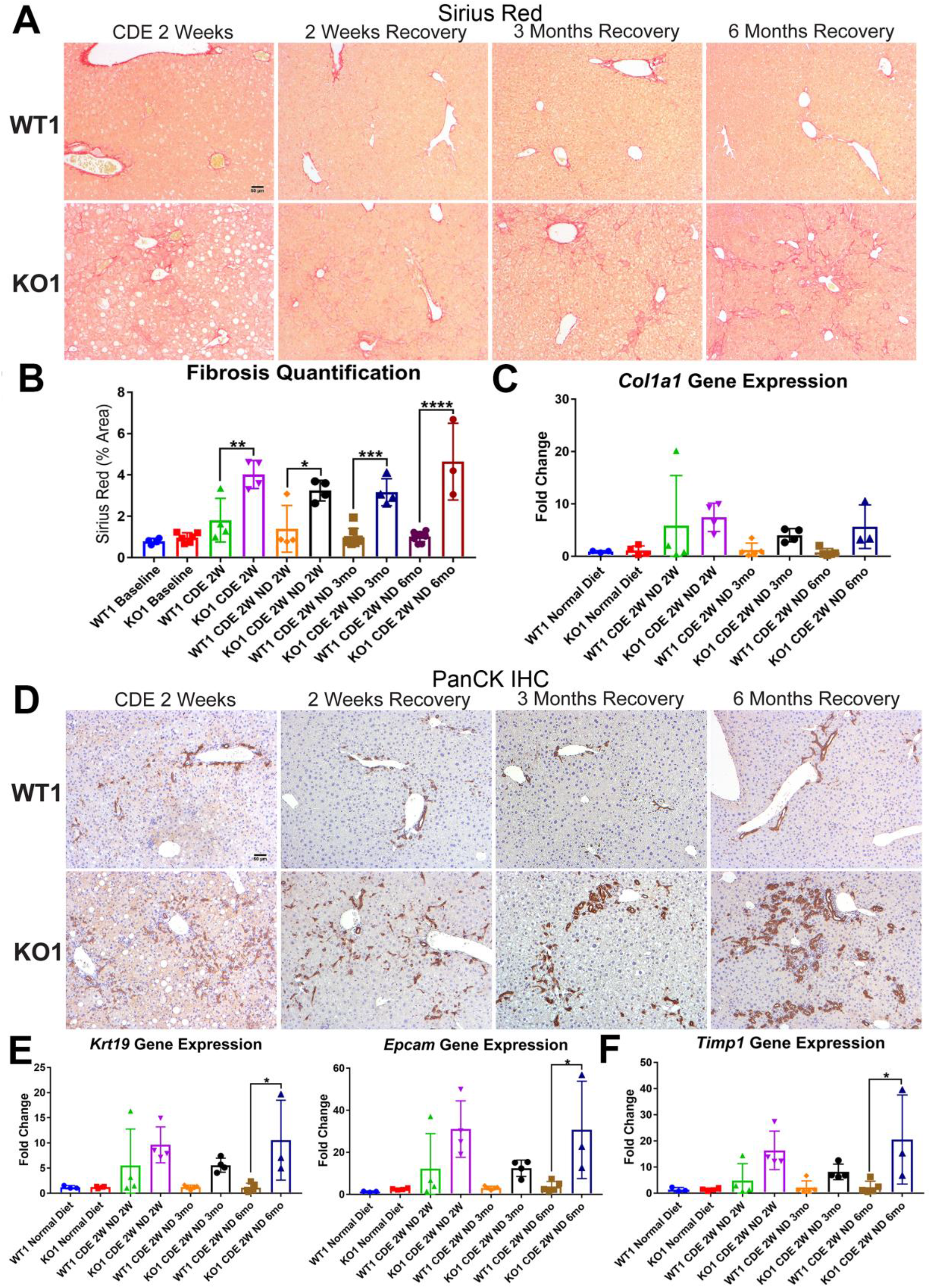
Unresolved fibrosis and ductular reaction in KO1 mice throughout 6m on recovery, after the initial 2w CDE diet injury. A) Sirius Red staining in WT1 and KO1 mice over time during recovery from CDE diet. Scale bar = 50µm. B) Quantification of Sirius Red staining (one way-ANOVA, *p<0.05, **p<0.01, ***p<0.001, ****p<0.0001. n = 3 to 6 per group. Individual animal values represented by dots.). C) A trend of increased expression of *Col1a1* in KO1 mice even at 6m of recovery (n = 3 to 5 per group. Individual animal values represented by dots.). D) PanCK staining in WT1 and KO1 mice over time during recovery from CDE diet. The DR changes from flattened, invasive and without lumen morphology from early time-points to numerous small luminal structures lined by a single layer of PanCK-positive columnar cells at 3m and 6m. Scale bar = 50µm. E) Significantly higher *Krt19* and *Epcam* gene expression in KO1 especially at 6m of recovery (one-way ANOVA, *p<0.05. n = 3 to 5 per group. Individual animal values represented by dots.). F) Significantly higher *Timp1* gene expression IN KO1 especially at 6m of recovery diet (one-way ANOVA, *p<0.05. n = 3 to 5 per group. Individual animal values represented by dots.)

DR was next assessed by IHC for PanCK. While there was a dramatic decrease in DR overtime in WT1 mice, a profound DR was observed in KO1 mice at all times, which was even more pronounced at 6m of recovery (Fig.4D). Furthermore, the DR was peculiar and composed of numerous small luminal structures lined by a single layer of PanCK-positive columnar cells at 3m and 6m, rather than more flattened and invasive DR without lumen seen at earlier stages of CDE injury and recovery in both WT1, KO1 and even KO2 (Fig.4D,2D). Enhanced gene expression for *Krt19* and *Epcam* was simultaneously evident in KO1 at these times (Fig.4E).

To determine if the continued DR was due to ongoing BEC proliferation, we co-stained KO1 livers from 3m recovery with PanCK and proliferating cell nuclear antigen (PCNA) (Fig.S1A). Significantly more BECs were proliferating in KO1 compared to WT1 mice (Fig. S1B). A subset of BECs in DR were also positive for phospho-Erk1/2 (p- Erk1/2), known for regulating BEC proliferation (Fig.S1C)^17^.

Since decreased gene expression of *Timp1* correlated with reduced fibrosis in KO2 at 6m of recovery, we next investigated its levels in KO1 and WT1. Timp1 tended to be upregulated in KO1 mice at all times but significantly at 6m recovery time (Fig.4F).

Taken together, these results suggest that KO1 mice which continue to lack β- catenin in hepithelial cells, show persistent Timp1 and fibrosis, and display continued and morphologically distinct DR associated with increased BEC proliferation at all times after the initial 2w CDE-diet injury, despite lack of active insult.

### Unremarkable changes in hepatic bile acids, apoptosis and senescence during resolution of fibrosis and DR during recovery from CDE diet in KO2 mice

To discern the basis of disparate DR and fibrosis between the two models, we first focused on investigating differences in specific injury processes between KO2 and WT2 during CDE injury and recovery. Increased hepatic bile acids have been implicated in hepatic injury and repair^18^ and our lab has previously reported altered hepatic bile acids (BAs) in KO1 mice after Methionine-Choline deficient (MCD) diet^19, 20^. However, CDE diet-fed KO2 or WT2 mice showed no significant increase in hepatic bile acids at any time suggesting these to not be driving DR or fibrosis in this model (Fig.S2A). We also investigated cell senescence as a possible driver of DR and fibrosis^21^. However, no significant hepatocyte senescence was observed by p21 IHC in WT2 or KO2 mice during long-term recovery from CDE diet (Fig.S2B).

Although ALTs were not elevated, we wanted to directly address any ongoing injury in recovering WT1 and KO1 mice. Cleaved caspase-3 staining showed minimal cell death in both WT1 and KO1 mice at 3m of recovery (Fig.S2C).

Thus, BA alterations, cellular senescence, and cell death are not the basis of DR and fibrosis in CDE injury, and hence can’t explain differences in recovery between KO1 and KO2 mice.

### Maintenance of adherens junctions in KO2 and KO1 during recovery from CDE- diet injury

Next, we assessed if continued absence of β-catenin in KO1 but not in KO2 at 6m of recovery could be affecting adherens junctions (AJs) integrity, and could explain differences in DR and fibrosis between KO2 and KO1 mice. Temporal disruption of cell-cell junctions has been associated with pathologies in hepatobiliary injury including after CDE diet^22^. However, at 6m of recovery, immunoprecipitation (IP) with E-cadherin showed an association of E-cadherin to β-catenin in KO2 and with γ-catenin in KO1, as has been shown in β-catenin-deficient livers by us previously (Fig.S2D)^23^.

Thus, intact AJs are present in both models during recovery, and can’t be the basis of sustained DR and fibrosis in KO1.

### Prolonged periportal inflammation in KO1 but not in KO2 mice during recovery from CDE diet correlates with ongoing DR and fibrosis

Previously, the presence of immune cell infiltration has been shown to be essential in development of DR and fibrosis. Mice lacking Th1 immune signaling or interferon-γ showed impaired DR and decreased fibrosis after CDE diet^24^. To address inflammation, we performed staining for CD45, a pan-leukocyte marker. There were high numbers of CD45-positive cells in WT2 and KO2 mice after 2w CDE diet, which declined in WT2 after 2w weeks of recovery but persisted pan-lobularly in KO2 up to 3m, and normalized to WT2 levels by 6m (Fig.5A). CD45-positive cells were present in high numbers in both WT1 and KO1 at 2w weeks of CDE-diet and while these numbers returned to baseline in WT2 at 2w of recovery, a more intense periportal appearing infiltration was seen in KO1 especially at 3m and 6m (Fig.5B).

**Figure 5:**
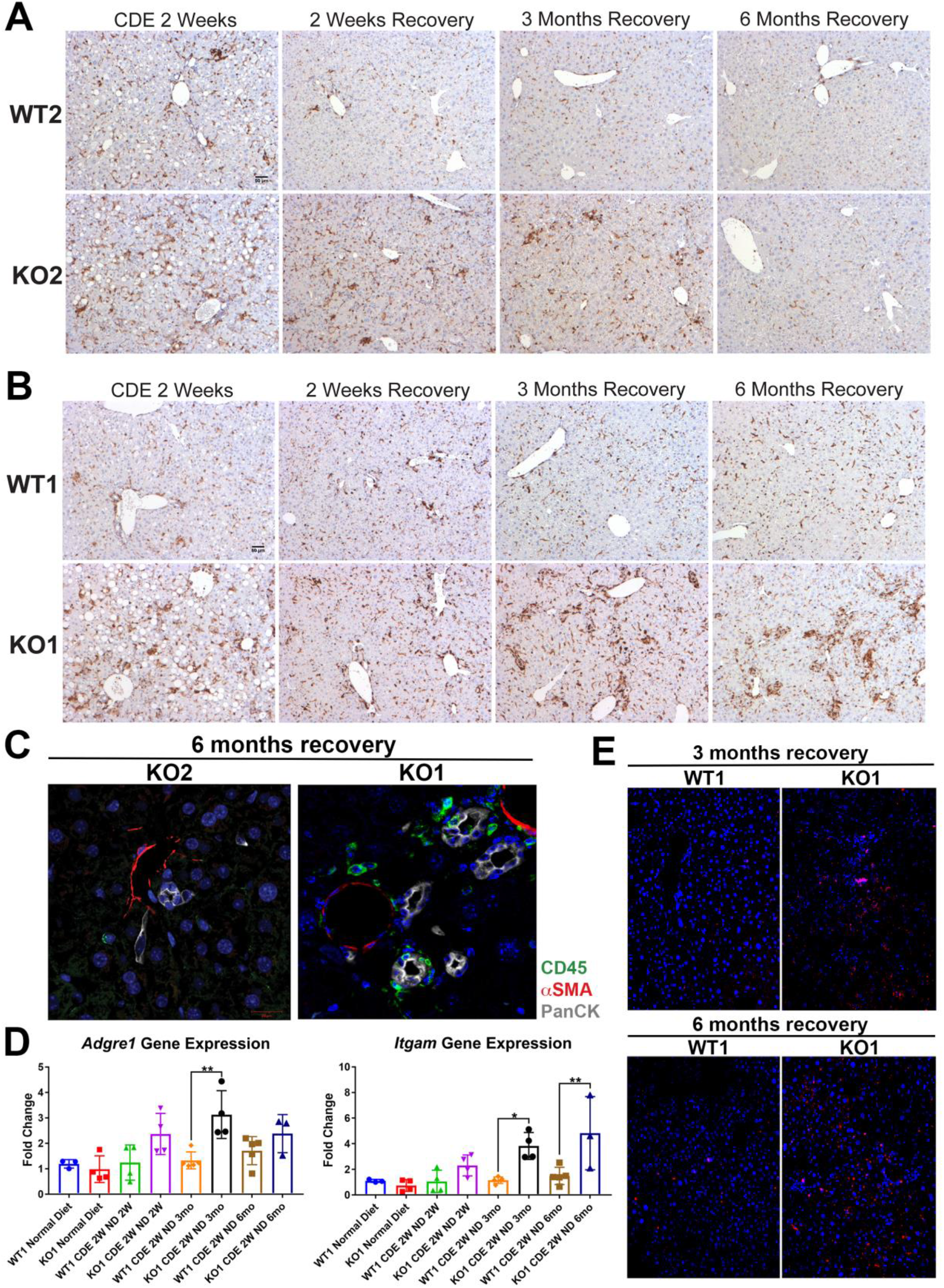
Sustained inflammation during recovery after the initial CDE diet injury in KO1 mice as compared to WT1, WT2 and KO2. A) CD45 immunostaining in WT2 and KO2 mice over time during recovery from CDE diet. Scale bar = 50µm. B) CD45 staining in WT1 and KO1 mice over time during recovery from CDE diet. Scale bar = 50µm. C) Representative confocal image of triple immunofluorescence for PanCK (white), αSMA (red), and CD45 (green) in KO1 and KO2 at 6m of recovery (400x). D) Changes in *Adgre1* and *Itgam* gene expression in WT1 versus KO1 at various time- points (one-way ANOVA, *p<0.05, **p<0.01. n = 3 to 5 per group. Individual animal values represented by dots.). E) Increased Ly6G staining (red) in KO1 mice after 3m and 6m of recovery (100x).

To determine if these inflammatory cells were close to DR, we performed triple immunofluorescence (IF) for PanCK, CD45, and myofibroblast marker α-smooth muscle actin (αSMA) (Fig.S3A). At 2w CDE diet, both WT2 and KO2 livers exhibited inflammatory cells and αSMA-positive cells close to BECs. After 2w of recovery, αSMA- positive cells were no longer associated with BECs in WT2 and no immune cells were seen. In KO2 mice, BECs, αSMA-positive cells and leukocytes were seen in close proximity to each other up to 3m months of recovery and returned to WT2-state at 6m (Fig.S3A). KO1 livers showed closely associated CD45- and PanCK-positive cells at all times in contrast to WT1 which lacked CD45 cells at all times after 2w of recovery (Fig.S3A). Intriguingly, even in KO1, αSMA-positive cells were not observed at any time after 2w of recovery. Overall, the presence of CD45-positive cells close to PanCK- positive cells was clear in KO1 but not in KO2 at 6m (Fig.5C).

Analysis in whole livers for expression of *Adgre1*, gene encoding macrophage marker F4/80, showed significant increase in KO1 compared to WT1 mice at 3m recovery (Fig.5D), and these macrophages were located close to the DR (Fig.S3B). Bone marrow monocyte-derived macrophages express high levels of *Itgam* (CD11b), and infiltrate liver during injury, express pro-inflammatory cytokines, and are involved in both progression and recovery phases of fibrosis^25^. *Itgam* expression was significantly higher in KO1 at 3m and 6m recovery times (Fig.5D). There was also increased staining for Ly6G, a marker for monocytes, granulocytes, and neutrophils, in KO1 at 3m and 6m recovery times (Fig.5E).

Together, these results show resolution of DR and fibrosis in KO2 correlated with reduced inflammation, whereas persistent β-catenin-negative DR and continuing fibrosis in KO1 mice at late recovery stages from CDE injury was associated with persistent periportal inflammation.

### Lack of β-catenin from BECs in KO1 leads to persistent NF-κB activation during recovery from CDE-diet induced injury

To address the mechanism of enhanced periportal inflammation, we next assessed the status of NF-κB, the master regulator of immune cell response. We have previously shown an inhibitory interaction of β-catenin with p65 subunit of NF-κB in hepatocytes and the absence of β-catenin in KO1 led to increased NF-κB activation in response to Lipopolysaccharide (LPS) or Tumor necrosis factor-α challenge^26^. Further, while immune cells are essential for DR and fibrosis, reactive ductules are a well-known source of pro-inflammatory and pro-fibrogenic cytokines and thus this cross-cellular signaling perpetuates overall injury^27, 28^. Since inflammatory cells were specifically enriched in KO1 in the periportal region and associated closely to the DR, we next assessed NF-κB status along with β-catenin in BECs in KO2 and KO1. At baseline in KO2, p65 subunit of NF-κB was present in the cytosol of the CK19-positive cells, as was β-catenin (Fig.6A). In KO1, at baseline, CK19-positive BECs lacked β-catenin and p65 was still evident in cytosol (Fig.6A). At 2w of CDE-diet, when DR and inflammation is ongoing in both KO2 and KO1 livers, a subset of CK19-positive BECs showed comparable nuclear p65 by confocal microscopy in both groups (Fig.6B,6C). At 6m, β-catenin-positive, CK19-positive BECs showed only cytosolic p65 similar to baseline (Fig.6B). However, unlike at baseline, KO1 livers at 6m recovery from CDE injury showed profound nuclear translocation of p65 in almost all CK19-positive BECs which continued to lack β-catenin (Fig.6B). Quantification showed significant difference in nuclear p65 in BECs in KO1 versus KO2 at 6m recovery time- point (Fig.6C).

**Figure 6:**
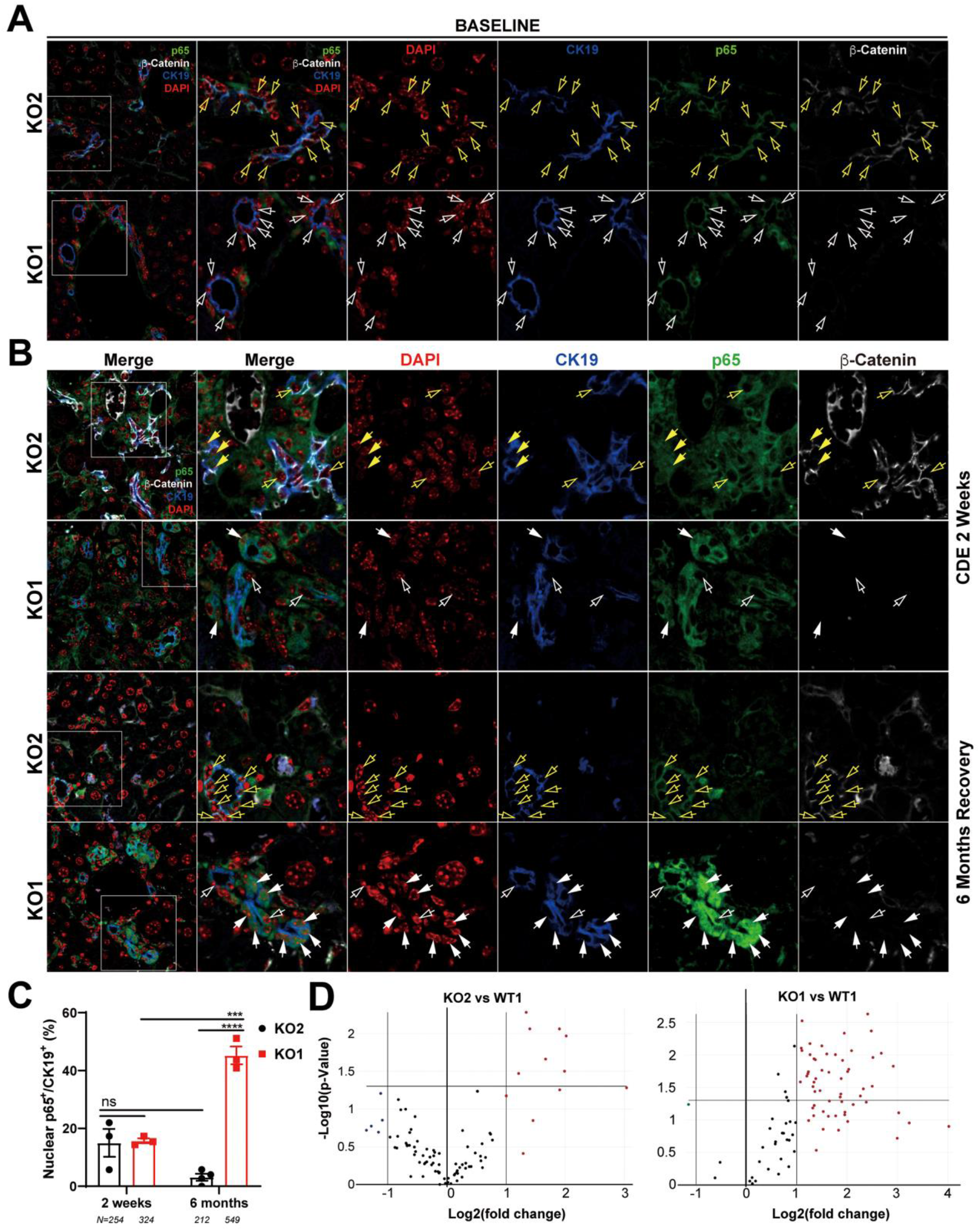
Nuclear translocation of p65 in BECs lacking β-catenin during recovery from CDE diet injury shows NF-κB activation in BECs only in KO1. A) Representative confocal image of triple immunofluorescence for CK19 (blue), β-catenin (white) and p65 (green) along with DAPI (red) in KO1 and KO2 baseline livers. Merged image at low magnification (100x) is shown in leftmost panel and higher magnification (200x) of selected area (box) along with its individual channels are shown to the right. Yellow open arrows identify CK19-positive BECs with cytosolic β-catenin and p65 in KO2. White open arrows identify CK19-positive BECs with cytosolic p65 and absent β- catenin in KO1. B) Representative confocal image of triple immunofluorescence for CK19 (blue), β-catenin (white) and p65 (green) along with DAPI (red) in KO1 and KO2 at 2w of CDE diet and after 6m of recovery on normal diet. The left most panel is low magnification (100x) merged image. The higher magnification (200x) of the selected boxed area is presented in the adjacent panel as a merged image followed by individual channels. Yellow open arrows identify CK19-positive BECs with cytosolic β-catenin and p65, and yellow solid arrows with nuclear p65 in KO2. White open arrows indicate CK19-positive BECs with cytosolic p65 and white solid arrows identify CK19-positive BECs with nuclear p65 with absent β-catenin in KO1. C) Quantification of CK19-positive cells showing nuclear p65 at 2w of CDE diet and 6m of recovery in KO2 versus KO1 (one way-ANOVA, ***p<0.001, ****p<0.0001. n = 3 per group. Individual animal values represented by dots. Number of cells counted are indicated.). D) Volcano plots of NF-κB downstream target gene expression in KO2, KO1 and WT1 livers at 6m of recovery. Genes with fold-change >2 are highlighted in red, with fold-change <2 are highlighted in green, and unchanged genes shown as black dots. (n = 3 per group)

To verify if nuclear p65 indicated NF-κB activation, 84 downstream target genes were checked by RT-PCR array (fold change threshold=>2, p-value threshold=0.05).

Relative to WT1, we found a striking upregulation in the expression of 44% of genes (37/84) in KO1, whereas 92% of genes (77/84) in KO2 livers were unchanged at 6m of recovery (Fig.6E). Clustergram showed KO2s were indistinguishable from WT1, but KO1 clearly separated from both groups (Fig.S4).

Altogether, these data suggest a pronounced and prolonged NF-κB activation in BECs lacking β-catenin along with periportal inflammation during recovery phase from CDE injury, whereas presence of β-catenin dampened NF-κB activation in BECs to curbed inflammation and assist in recovery from the same injury.

### β-Catenin modulation in BECs leads to differential impact on NF-κB activity through β-catenin and p65 interactions

To more conclusively address the relationship between β-catenin and NF-κB in BECs directly, we utilized immortalized mouse small cholangiocyte cells (SMCCs), which were transfected with control- or β- catenin siRNA together with either β-catenin-TCF Topflash reporter or p65 luciferase reporter. Knockdown of β-catenin in SMCCs shown by a significant decrease in Topflash activity, induced p65 luciferase activity, which was 3-times greater than caused by 100ng/ml LPS stimulation (Fig.7A). Conversely, expression of stable S45Y-β-catenin or T41A-β-catenin (not shown) in SMCCs led to increased Topflash activity but significantly decreased p65 reporter activity with or without LPS (Fig.7B). Similar negative regulation was also observed in MzChA and HuCCT1, two independent human BEC lines (Fig.S5A,5B).

**Figure 7:**
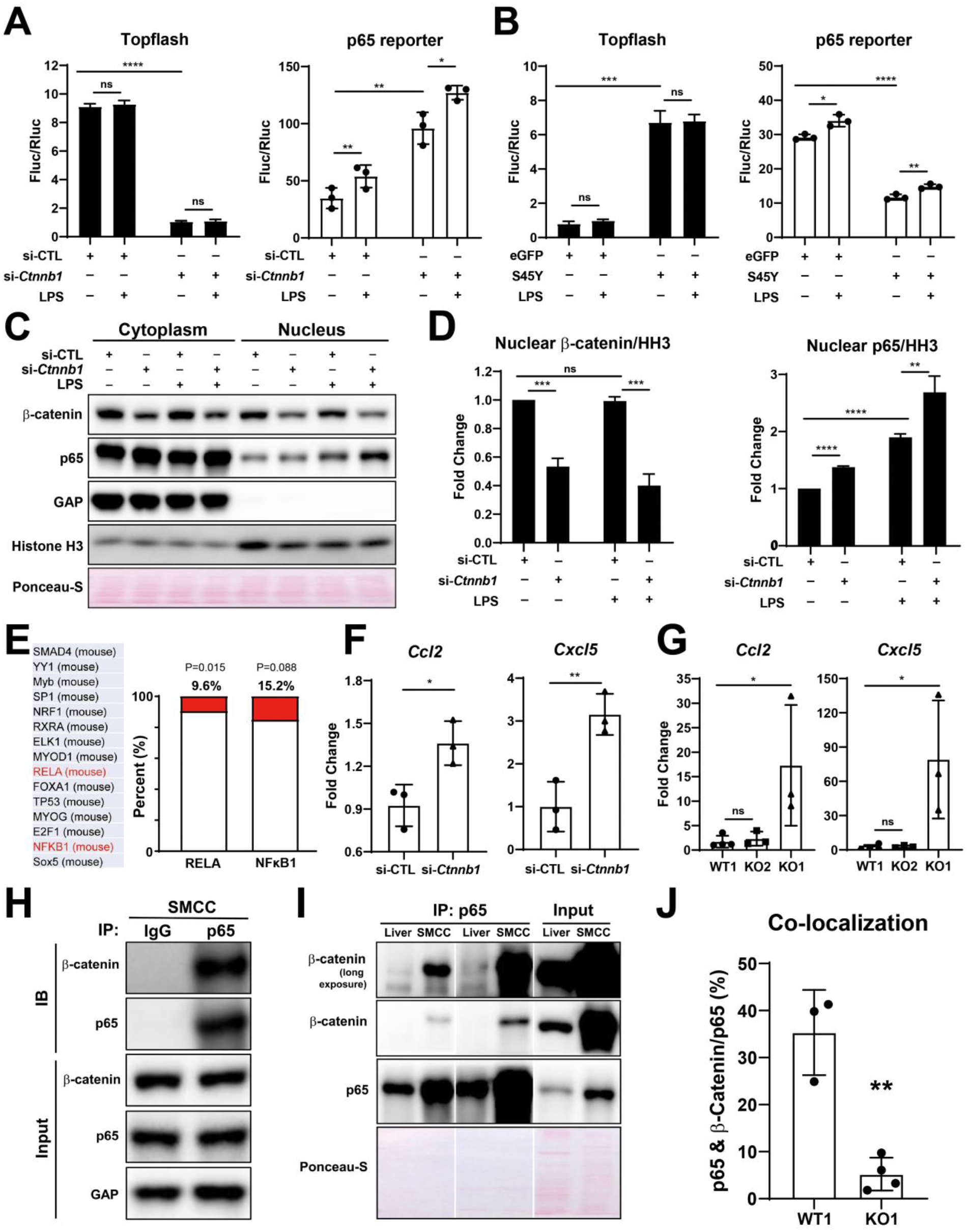
Modulation of β-catenin in BECs perturbs its complex with p65 to impact NF-κB activity. A) Luciferase reporter assay shows successful knockdown of *Ctnnb1* in SMCC line by Topflash assay (left), which stimulates p65 transcriptional activity with or without 100ng/ml LPS (right) (Unpaired t-test, ns: no significance, *p<0.05, **p<0.01, ****p<0.0001. n = 3 biological replication). B) Luciferase reporter assay shows expression of constitutively active S45Y-β-catenin enhances Topflash (left) and suppresses p65 transcriptional activity with or without 100ng/ml LPS (right) (Unpaired t- test, ns: no significance, *p<0.05, **p<0.01, ***p<0.001, ****p<0.0001, n = 3 biological replication). C) Representative WB from two independent experiment shows knockdown of *Ctnnb1* increases p65 nuclear translocation with or without 500ng/ml LPS. D) Quantification of nuclear β-catenin (left) and nuclear p65 (right) to HH3 (Blots in Figure 7C were technically quantified three times and p-value was calculated using unpaired t- test, ns: no significance, **p<0.01, ***p<0.001, ****p<0.0001). E) Identification of RELA and NFKB1 among the top fifteen transcription factors identified by applying the 335 DEGs to JASPAR. F) qPCR shows knockdown of *Ctnnb1* in SMCCs induces *Ccl2* (left) and *Cxcl5* (right) expression (Unpaired t-test, ns: no significance, *p<0.05, **p<0.01, n = 3 biological replication). G) qPCR shows *Ccl2* (left) and *Cxcl5* (right) are induced in KO1 after 6m recovery of CDE diet (Unpaired t-test, *p<0.05. n = 3 to 4 biological replication). H) Representative Immunoprecipitation (IP) image from two independent experiment shows p65 is strongly associated with β-catenin in SMCC. I) IP shows that p65 is associated with β-catenin in whole liver lysate (L: Liver; S: SMCC; P: equal amount of liver and SMCC lysate). J) Quantification of colocalization of p65 and β-catenin is significantly diminished in KO1 compare to WT1 (Unpaired t-test, **p<0.01, n = 3 to 4 biological replication).

To address how β-catenin regulates NF-κB activity, SMCCs transfected with control- or Ctnnb1-siRNA were cultured with or without LPS, and subjected to cell fractionation. WB using nuclear- and cytoplasmic-enriched fractions showed significantly higher levels of p65 in the nuclear compartment in β-catenin-silenced group versus controls, both with or without LPS (Fig.7C,7D).

Next, we modulated β-catenin activity in SMCCs to determine changes in global gene expression. Bulk RNA-seq was performed on β-catenin-silenced or control SMCCs, and on SMCCs transfected with S45Y-β-catenin or eGFP. Using a cutoff p-value≤0.05 and abs(log2FC)≥1.5, we found 335 β-catenin-regulated genes in BECs. Specifically, β-Catenin-silenced SMCCs showed 75 upregulated and 122 downregulated, and β-catenin-active SMCCs showed 76 upregulated and 69 downregulated genes (Fig.S5C,5D). While there was a minimal overlap (Fig.S5E), JASPAR was queried to identify transcription factor (TF) binding profiles in the 335 DEGs. RELA (p65) was identified among the top TFs (ranking by p-value), with 9.6% of DEGs showing known RELA regulation (32/335, p=0.015), and 15.2% (51/335, p=0.088) showing NF-κB binding sites (Fig.7E). Interestingly, from RNA-seq and by qPCR, we found modest but significant increase in CCL2, and a more pronounced and significant increase in CXCL5 expression, in the β-catenin-silenced SMCCs (Fig.7F).

After 6m recovery from CDE diet, a significant induction in CCL2 and CXCL5 expression was noted in KO1 but not in WT1 and KO2 (Fig.7G).

Next, we investigated if β-catenin interacts with p65 in BECs. Whole cell lysates from SMCCs and normal mouse liver were used to pulldown p65. We identified robust β-catenin association with p65 in SMCCs (Fig.7H,7I). A fainter but positive β-catenin- p65 association was evident in whole livers, likely due to low BEC representation in protein lysates from whole livers (Fig.7I). To further verify presence of β-catenin-p65 complex in BECs *in vivo*, we examined β-catenin and p65 localization using confocal microscopy. In WT1 liver at baseline, a notable co-localization of p65 and β-catenin was evident in BECs which was absent in KO1 (Fig.7J,S5F). Quantification of co-localization using Image J showed about 35% of p65 is associated with β-catenin, which was significantly greater than KO1 (Fig.7J).

Altogether, biochemical and IF studies identify a heretofore undescribed β- catenin-p65 complex in BECs, Further, β-catenin seems to be important in shutting off NF-κB activation when its signaling is not required anymore and absence of β-catenin in BECs leads to sustained and excessive NF-κB activation along with its associated sequela- proliferation, inflammation and fibrosis^29, 30^.

### Nuclear p65 in pathologic DR in subset of clinical cases divulges heretofore unidentified interactions of β-catenin, p65 and CFTR

Since persistent DR observed in KO1 at 3m and 6m after recovery from CDE injury showed unique morphology consisting of numerous small luminal structures lined by a single layer of PanCK- positive cells, we decided to interrogate livers from clinical cases exhibiting DR including alcoholic hepatitis (AH), polycystic liver disease (PLD) and cystic fibrosis (CF). As seen by panCK IF, AH cases displayed DR with variable morphology including areas of luminal small DR (shown in representative case), whereas PLD showed large cysts lined by flattened BECs (Fig.8A). DR in CF was more homogeneous and appeared uniformly reminiscent of what we observed in 3m and 6m recovery times in KO1 (Fig.4D, 8A). Intriguingly, strongest and significant nuclear p65 was consistently observed in DR seen in CF cases followed by PLD with only a very small subset of cells in DR showing nuclear p65 in AH (Fig.8B). While β-catenin seem to be unaltered by IF staining in all three pathologies (Fig.8A), we observed a decrease in total β-catenin in a single CF case from whom two independent frozen liver samples were available (Fig.8C).

**Figure 8.**
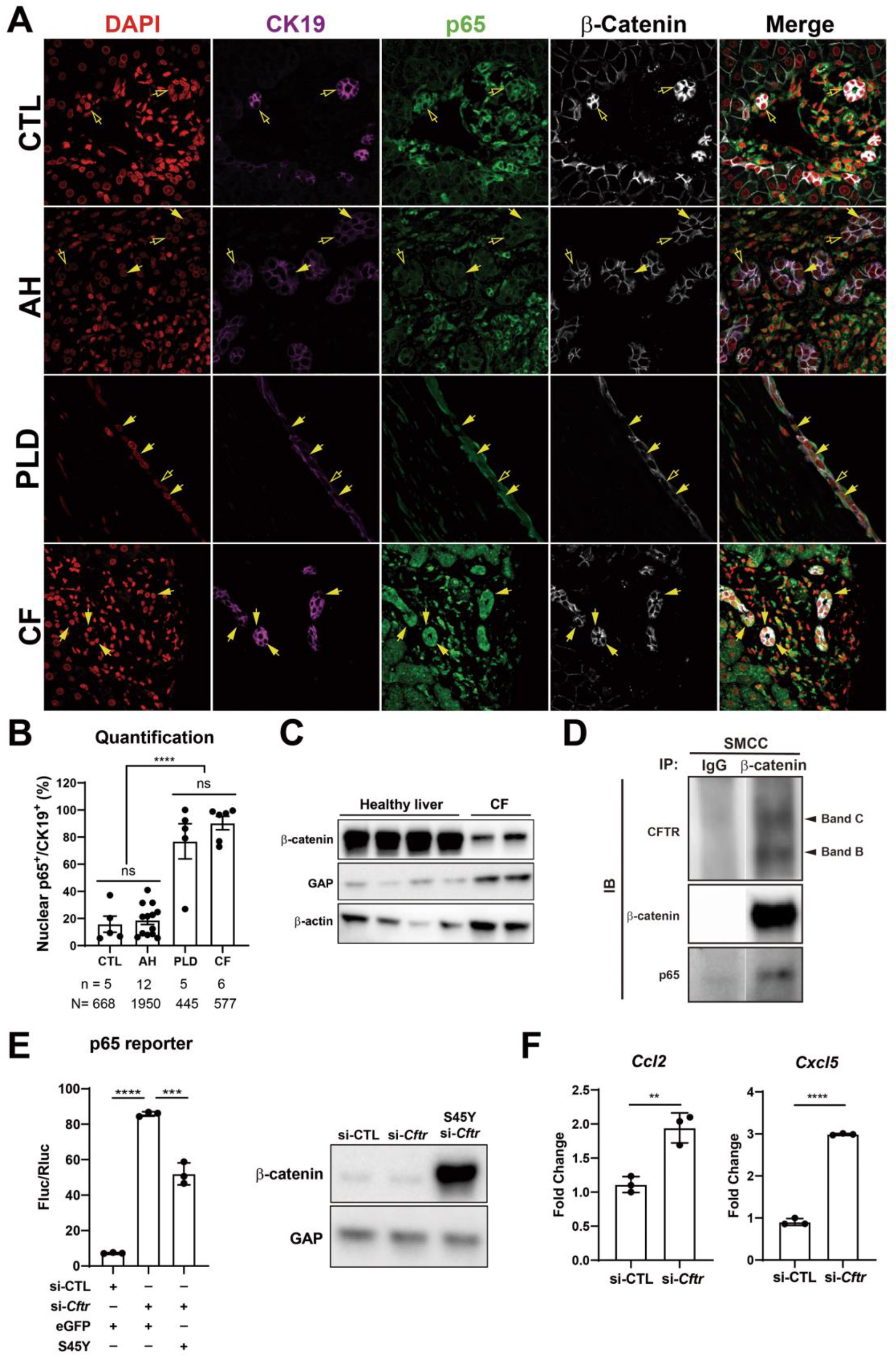
Nuclear p65 is highly present in the ductular cells of cystic type liver disease. A) Representative IF images of liver sections from patients with healthy liver (CTL), alcoholic hepatitis (AH), polycystic liver disease (PLD), and cystic fibrosis (CF). (200x) B) Quantification of the percentage of nuclear p65+CK19+ cells among CK19+ BECs from CTL (668 cells from 5 patients), AH (1950 cells from 12 patients), PF (445 cells from 5 patients), and CF (577 cells from 6 patients) (one-way ANOVA, ****p<0.0001). C) WB shows β-catenin is decreased in two liver samples from one CF patient as compared to four healthy liver controls. D) IP studies show CFTR (C-Band and B-Band) and p65 to be pulled down with β-catenin and not with IgG control in SMCC. Images are from the same gel with same exposure time. E) Luciferase reporter assay shows knockdown of CFTR in SMCC line strongly induces p65 transcriptional activity. Overexpression of stable S45Y-β-catenin partially rescues p65 activity (Unpaired t-test, ***p<0.001, ****p<0.0001, n = 3 biological replication). Representative WB shows β-catenin levels after si-*Cftr* or after simultaneous S45Y-β-catenin expression as compared to si-control (si-CTL). F) qPCR shows knockdown of CFTR induces *Ccl2* and *Cxcl5* expression in SMCC (Unpaired t-test, **p<0.01, ***p<0.001, n = 3 biological replication).

CF cases typically have varying loss-of-function (LOF) mutations in *CFTR* gene. Since β-catenin-p65 interactions were observed in SMCCs, we next assessed if CFTR is interacting with this complex. We observed concomitant pulldown of both B-Band (faster migrating core glycosylated immature form) as well as slowly migrating C-Band (complex glycosylated form)^31^, along with p65, when we immunoprecipitated β-catenin in SMCCs (Fig.8D). To mimic LOF of CFTR seen in CF, we next silenced *Cftr* in SMCCs. Knockdown of *Cftr* led to a pronounced increase in p65 reporter activity (Fig.8E). Likewise, the expression of NF-κB target chemokines *Ccl2* and *Cxcl5,* were significantly induced upon *Cftr* knockdown (Fig.8F). To query impact of β-catenin modulation, we next co-expressed stable-β-catenin (S45Y-mutant) in control and *Cftr-* siRNA-transfected SMCCs. Stabilization of β-catenin significantly decreased CFTR knockdown-induced p65 activation (Fig.8E).

Thus, we identify important interactions between β-catenin, p65 and CFTR in BECs, and LOF of CFTR or β-catenin leads to enhanced p65 activation, which can be inhibited by β-catenin stabilization.

## Discussion

DR is a common hallmark of many chronic liver pathologies although it’s morphology is heterogeneous ranging from isolated invasive ductular cells, luminal phenotype and sometimes purely cystic^5, 6, 32^. The significance of DR remains controversial and has been associated with both repair and disease progression^5, 33^. Its role as a source of *de novo* hepatocytes through the process of transdifferentiation is indisputable in preclinical models shown by many fate-tracing studies^14, 21, 32, 34^. At the same time, DR can induce fibrosis by secreting pro-inflammatory and profibrogenic factors to contribute to the disease process^7,8,9, 29, 35^. What drives the pro-inflammatory and pro-fibrogenic phenotype of these reactive BECs and what reverts these cells back to normal, is poorly understood.

Previously, we and others have described β-catenin-p65 complex in hepatocytes, breast and colon cancer cells, which could inhibit NF-κB activation^26, 36^. The exact biological significance of this interaction is not well understood, and likely context dependent. We identify this complex in normal cholangiocytes demonstrating its critical role in homeostasis in this cell-type. Being a ‘sticky’ protein, β-catenin interacts with many proteins in a cell to modulate their activities^37^. We show that β-catenin-p65 complex, keeps NF-κB activity in check by preventing its nuclear translocation. β- Catenin activation is observed in BECs in DR during chronic liver injuries^38,39,40^. Unlike in hepatocytes, β-catenin activation in BECs does not play a role in proliferation^14, 40^.

Our current study shows β-catenin stabilization in BECs may in fact be ‘mopping’ up p65 to dampen and eventually shutting off NF-κB activation, reverting BECs to their quiescence. NF-κB activation has been shown to be important in BEC proliferation and DR by regulating Jagged/Notch signaling^29^. And NF-κB in BECs has been suggested to play a role in inducing profibrogenic and pro-inflammatory milieu as well^41^. Absence of β-catenin in BECs in KO1 prevented formation of a complex with p65, allowing unchecked NF-κB activation whereas re-formation of β-catenin-p65 complex in KO2 as a way of keeping inflammation in check during chronic injuries. Molecular underpinnings of β-catenin stabilization in BECs during liver injuries remains under investigation although portal fibroblasts, macrophages, hepatocytes and BECs have all been shown to secrete ligands like Wnt7a, Wnt7b, Wnt10a and Wnt5a^39, 40, 42^.

Another intriguing observation was the distinct morphology of DR evident in the recovery phase from the CDE injury in β-catenin-deficient livers, which was reminiscent histology in CF cases^43^. Surprisingly, very few BECs in AH showed nuclear p65, while PLD cases showed variable but increased nuclear p65 in cells lining the cysts.

However, DR in CF cases exhibited strong and consistent nuclear p65. This led us to investigate interactions between β-catenin-p65 and CFTR, whose gene is mutated in CF. CFTR protein is present in BECs only in the liver^44^. We observed a pulldown of CFTR (B-Band and C-Band) with β-catenin in BEC line. To mimic LOF, which is the common end-result of *CFTR* mutations in CF patients, we silenced CFTR in SMCC line, which led to a profound p65 activation that was decreased upon β-catenin stabilization. This suggests an important tripartite regulatory interaction between these proteins.

Interestingly, in human lung epithelial cells, a proteomic screen identified interaction of β-catenin with WT CFTR but not with ΔF508 CFTR, a major site of LOF mutation in the *CFTR*^45^. Additionally, in mouse intestine, CFTR was shown to stabilize β-catenin and prevent its degradation^46^. The same study showed that ΔF508 CFTR is unable to interact with β-catenin leading to β-catenin degradation and eventually resulting in activation of NF-κB-mediated inflammatory cascade. Our results show an existence of a tripartite interaction between β-catenin, p65 and CFTR in cholangiocytes and LOF of CFTR led to reduced β-catenin protein in BECs both *in vitro* (SMCC) and *in vivo* (CF patient liver lysate), leading to profound NF-κB activation and increased expression of its pro-inflammatory chemokine targets. We believe that the classical pathology of liver disease in CF including periductal inflammation, DR, periportal fibrosis, and focal biliary cirrhosis, may be explained by our observations, in addition to previously described mechanisms such as Rous sarcoma oncogene cellular homolog (Src) dependent toll like receptor-4 activation^47^. Since β-catenin activation in BECs inhibited NF-κB activation due to CFTR-silencing, this strategy may have therapeutic implications in controlling CF disease progression in the liver and elsewhere and future studies will directly address this novelty. NF-κB activation has been shown to be important in BEC proliferation and DR by regulating Jagged/Notch signaling^29^. The same study showed NF-κB activation to be regulated by Cystein-rich protein 61 (CYR61), whose knockdown reduced DR. Incidentally, *CFTR* was also significantly reduced in that study and might have been a mechanism of DR through disruption of CTFR-p65-β-catenin interactions.

The mechanism of NF-κB activation and whether these tripartite interactions are playing any role in other pathologies such PLD or subset of AH cases, requires further investigation.

There is very little understanding of the process of resolution of DR along with its associated fibrosis, although these are strongly linked to inflammation^48^. Our study provides novel insight into not only the molecular underpinnings of a reactive cholangiocyte, but also sheds light on how BECs regulate the immune microenvironment. Levels of Timp1 correlate with fibrosis, and several groups have investigated the use of TIMP1 levels as a biomarker for fibrosis in hepatitis C patients^49, 50^. Mice with overexpression of TIMP1 developed dramatically more fibrosis after CCl_4_ treatment^15^, and TIMP1 transgenic mice showed impaired fibrosis resolution after cessation of CCl_4_^16^. Our data is consistent since expression of *Timp1* was associated with fibrosis, and levels of *Timp1* normalized upon resolution of fibrosis.

## Materials and Methods

### Animals

All animals are housed in temperature and light-controlled facilities and are maintained in accordance with the Guide for Care and Use of Laboratory Animals and the Animal Welfare Act. Generation of *Albumin-Cre;Ctnnb1^flox/flox^* mice and wild-type littermates has been described previously^12^. Generation of *Ctnnb1^flox/flox^*;*Rosa-stop^flox/flox^- EYFP* reporter mice was also described previously^14^. In brief, 23–25 day-old *Ctnnb1^flox/flox^*;*Rosa-stop^flox/flox^-EYFP* mice were injected intraperitoneally with 1x10^12^ genome copies (GCs) of adeno-associated virus serotype 8 encoding Cre recombinase under the hepatocyte-specific thyroid binding globulin promoter (AAV8-TBG-Cre) (Addgene) followed by a 12 days (12d) washout period. The same protocol was utilized on *Ctnnb1^+/+^*;*Rosa-stop^flox/flox^-EYFP* mice to generate WT2 mice. When mice were 4-6 weeks old, choline-deficient diet (Envigo Teklad Diets) supplemented with 0.15% ethionine drinking water (Acros Organics, 146170100) was administered for 2 weeks.

For recovery time points, animals were switched back to normal chow diet for up to 6 months. Serum biochemistry analysis was performed by automated methods at the University of Pittsburgh Medical Center clinical chemistry laboratory. All studies were performed according to the guidelines of the National Institutes of Health and the University of Pittsburgh Institutional Animal Use and Care Committee.

### Patient data

All patient tissue sections were provided by the Pittsburgh Liver Research Center’s (PLRC’s) Clinical Biospecimen Repository and Processing Core (CBPRC), supported by P30DK120531. Sections from 5 patients with healthy liver, 12 patients with DR from AH (n=10) and/or NASH (n=2), 5 patients with DR associated with PLD, and 6 patients with DR in CF cases were triple stained with CK19, p65, β-catenin for further analysis. Patient information from these groups of cases is listed in Supplementary Table 1. Two pieces of frozen livers from one CF patient (TP10-P531) was provided by Pitt Biospecimen Core and used for WB. Information on this case is also included in Supplementary Table 1.

### Immunohistochemistry

The IHC protocols have been described previously^14^. In brief, liver tissue was fixed in 10% buffered formalin for 48 hours prior to paraffin embedding. Blocks were cut into 4µm sections, deparaffinized, and washed with PBS. For antigen retrieval, samples were microwaved for 12 minutes in pH 6 sodium citrate buffer (PanCK, CD45, p-Erk1/2, Cleaved Caspase 3) or Tris-EDTA buffer (p21), were pressure cooked for 20 minutes in pH 6 sodium citrate buffer (β-catenin), or were incubated with Proteinase K (Agilent Dako, S302030-2) for 10 minutes (F4/80).

Samples were then placed in 3% H_2_O_2_ for 10 minutes to quench endogenous peroxide activity. After washing with PBS, slides were blocked with Super Block (ScyTek Laboratories, AAA500) for 10 minutes or 10% goat serum in PBS for 10 minutes (p21).

The primary antibodies were incubated at the following concentrations in antibody diluent (PBS + 1% BSA (Fisher BioReagents, BP1605-100) with 0.1% Tween™ 20 (Fisher BioReagents, BP337-500): PanCK (Dako, Z0622, 1:200), Cleaved Caspase 3 (Cell Signaling, 9664, 1:100), p-Erk1/2 (Cell Signaling, 4370, 1:100), F4/80 (BioRad, MCA497A488, 1:100) for one hour at room temperature or at 4°C overnight: β-catenin (Abcam, ab32572, 1:50) and p21 (Santa Cruz, sc-471, 1:25). Samples were washed with PBS three times and incubated with the appropriate biotinylated secondary antibody (Vector Laboratories) diluted 1:500 in antibody diluent for 30 minutes at room temperature. Samples were washed with PBS three times and sensitized with the Vectastain® ABC kit (Vector Laboratories, PK-6101) for 30 minutes. Following three washes with PBS color was developed with DAB Peroxidase Substrate Kit (Vector Laboratories, SK-4100), followed by quenching in distilled water for five minutes. Slides were counterstained with hematoxylin (Thermo Scientific, 7211), dehydrated to xylene and coverslips applied with Cytoseal™ XYL (Thermo Scientific, 8312-4). For Sirius Red staining, samples were deparaffinized and incubated for one hour in Picro-Sirius Red Stain (American MasterTech, STPSRPT), washed twice in 0.5% acetic acid water, dehydrated to xylene, and coverslipped. Images were taken on a Zeiss Axioskop 40 inverted brightfield microscope.

### Immunofluorescence

Liver tissue was fixed in 10% buffered formalin overnight, cryopreserved with 30% sucrose in PBS overnight, frozen in OCT compound (Sakura, 4583) and stored at -80°C. OCT-embedded samples were cut into 5 µm sections, allowed to air-dry, and then washed in PBS. Antigen retrieval was performed through microwaving in pH 6 sodium citrate buffer. Slides were washed with PBS and permeabilized with 0.1% Triton X-100 in PBS for 20 minutes at room temperature. Samples were washed three times with PBS and then blocked with 2% Donkey serum in 0.1% Tween™ 20 in PBS (antibody diluent) for 30 minutes at room temperature.

Antibodies were diluted as follows: PanCK (Dako Z0622, 1:200), PCNA (Santa Cruz Biotechnology, sc-56, 1:1000), p65 (Santa Cruz Biotechnology, sc-372, 1:500), β- catenin (BD Biosciences, 610154, 1:500), CK-19 (DSHB, TROMA-III, 1:10) in antibody diluent and incubated at 4°C overnight. Ly6G (Clone: RB6-8C5) antibody was purchased from Thermo Fisher Scientific, Waltham, MA. Samples were washed three times in PBS and incubated with the proper fluorescent secondary antibody (AlexaFluor 488/555/647, Invitrogen) diluted 1:400 in antibody diluent for two hours at room temperature. The αSMA antibody is directly conjugated to Cy3 and requires no secondary antibody. Samples were washed three times with PBS and incubated with DAPI (Sigma, B2883) for 1 minute. Samples were washed three times with PBS and mounted with fluoromount (SouthernBiotech) or ProLong^TM^ Gold antifade reagent (Invitrogen, P10144). Images were taken on a Nikon Eclipse Ti epifluorescence microscope or a Zeiss LSM700 confocal microscope.

### Image quantification

To determine BEC proliferation, for each sample seven images at x200 magnification of periportal regions were taken and in each image the number of PanCK+/PCNA+ cells was manually counted. For quantification of Sirius Red staining, staining intensity was measured in ImageJ for five images at 100x magnification per sample. To determine p65 nuclear positive cholangiocytes, four images from each animal or patient at x200 magnification of ductular reaction regions were counted.

### RT-PCR

Whole liver was homogenized in TRIzol™ (Thermo Scientific, 15596026) and nucleic acid was isolated through phenol-chloroform extraction. Cellular DNA was digested with DNA-free™ Kit (ambion, AM1906), and RNA was reverse-transcribed into cDNA using SuperScript® III (Invitrogen, 18080-044). Real-time PCR was performed in technical triplicate on a StepOnePlus™ Real-Time PCR System (Applied Biosystems, 4376600) or on a BioRad CFX96 Real-Time System using the Power SYBR® Green PCR Master Mix (Applied Biosystems, 4367660). Target gene expression was normalized to the average of two housekeeping genes (*Gapdh* and *Rn18s*), and fold change was calculated utilizing the ΔΔ-Ct method. Primers are listed in Supplementary Table 2. For RT-PCR Arrays, 2μl of cDNA, 7.5μl of Power SYBR® Green PCR Master Mix and 4.5μl of nuclease-free water, were premixed and added to each well of RT² Profiler™ PCR Array Mouse NFκB Signaling Pathway (Qiagen, PAMM-025Z) for target gene qPCR. RT-PCR arrays data were analyzed at GeneGlobe (https://geneglobe.qiagen.com/us/analyze/). The average of three housekeeping genes (*Rn18s, Actb* and *Gapdh*) was used for normalization. Volcano plot and clustergram were generated by the data analysis web portal mentioned above.

### Immunoprecipitation and Western blot

Whole liver tissue was homogenized in RIPA buffer premixed with fresh protease and phosphatase inhibitor cocktails. Cytoplasmic and Nuclear extracts were prepared using the NE-PER^TM^ Nuclear and Cytoplasmic Extraction Reagents (Thermo Fisher Scientific, 78835). The concentration of the protein was determined by the bicinchoninic acid assay. For immunoprecipitation, 1mg of SMCC lysate was precleared with 40μl of Protein A/G PLUS-Agarose (Santa Cruz Biotechnology, sc-2003) for 2 hours at 4°C. After centrifugation (3,000 rpm, 1 minute), the supernatant was incubated with 2μg of p65 antibody (Santa Cruz Biotechnology, sc- 8008, 1:100), 2μg of β-catenin antibody (BD Biosciences, 610154, 1:100) or control IgG overnight at 4°C. The next day, samples were incubated with 40μl of Protein A/G PLUS-Agarose for 1 hour at 4°C. The pellet was collected, washed with RIPA buffer for 3 times, resuspended in 10μl of loading buffer, and subjected to electrophoresis. Protein lysate was separated on pre-cast 7.5% or 4-20% polyacrylamide gels (Bio-Rad) and transferred to the PVDF membrane using the Trans-Blot Turbo Transfer System (Bio-Rad). Membranes were blocked for 75 minutes with 5% skim milk (Lab Scientific, M0841) or 5% BSA in Blotto buffer (0.15M NaCl, 0.02M Tris pH 7.5, 0.1% Tween in dH2O), and incubated with primary antibodies at 4°C overnight at the following concentrations: β-catenin (Cell Signaling, 8480, 1:1000 in 1% BSA), p65 (Cell Signaling, 8242, 1:1000 in 1% BSA), CFTR (Alomone labs, ACL-006, 1:250 in 1% BSA), GAPDH (Cell Signaling, 5174, 1:10000 in 1% milk), Histone H3 (Cell Signaling, 9715, 1:1000 in 1% milk). Membranes were washed in Blotto buffer and incubated with the appropriate HRP-conjugated secondary antibody for 75 minutes at room temperature. Membranes were washed with Blotto buffer, and bands were developed utilizing SuperSignal® West Pico Chemiluminescent Substrate (Thermo Scientific, 34080) and visualized by autoradiography.

### Cell culture and reporter assays

SMCCs, MzChA and HuCCT1 were seeded on 6 well plates in a humidity-saturated incubator with 5% CO2 maintained at 37°C. For p65 reporter assay, Cells were transfected with 2μg of p65 reporter and 0.2μg of *Renilla* (internal control) together with 2μg of either eGFP (control) or S45Y to overexpress constitutively active β-catenin using Lipofectamine™ 3000 Transfection Reagent (Invitrogen™, L3000008), or together with si-Control (Cell Signaling, 6568) and si-β- catenin (Cell Signaling, 6225) to knockdown β-catenin using Lipofectamine™ RNAiMAX Transfection Reagent (Invitrogen™, 13778150). For TopFlash reporter assay, p65 reporter above was replaced by TopFlash plasmid. Cells were treated with 100ng/ml of LPS 6 hours before harvest. si-CFTR (Santa Cruz Biotech, sc-35053) was used to knockdown CFTR in SMCCs. Cells were harvested at 48 hours and luciferase signals were got using Dual-Luciferase® Reporter Assay System (Progema, E1910) and normalized to the value of *Renilla*.

### Measurement of hepatic bile acids

Total hepatic bile acids were measured using the Mouse Total Bile Acids Assay Kit from Crystal Chem (Downers Grove, IL), as per the manufacturer’s instructions. To isolate total bile acids from liver, 50-100mg frozen liver tissue was homogenized in 70% ethanol at room temperature, then samples were incubated in capped glass tubes at 50°C for 2 hours. The homogenates were centrifuged at 6,000g for 10 minutes to collect the supernatant. Total bile acid concentrations were determined using the calibration curve from the standard provided in the kit and the mean change in absorbance value for each sample.

### RNAseq and analysis

Twelve SMCC RNA samples were measured: 3 for CTNNB1 activation (SMCC-S45Y), 3 for CTNNB1 activation control (SMCC-eGFP), 3 for CTNNB1 silencing (SMCC-si-β-catenin), 3 for CTNNB1 silencing control (SMCC-si- Control). In total 12 RNA-seq libraries were sequenced. For each library, quality control was performed to each raw sequencing data by tool FastQC. Based on the QC results, low-quality reads and adapter sequences were filtered out by tool Trimmomatic^51^.

Surviving reads were then aligned to mouse reference genome mm10 by aligner Hisat2^52^. HTSeq tool^53^ was then applied to the aligned file for gene quantification. Based on the gene count, differential expression analysis was applied to compare CTNNB1 activation samples with their corresponding controls, and to compare CTNNB1 silencing samples with their corresponding controls, respectively. R package DESeq2^54^ was employed to perform the test and differentially expressed genes (DEGs) were defined as genes with fold-change higher than 1.5-fold and p-value (or adjusted p-value) smaller than 0.05. These DEGs were further used to detect common upstream transcription factors based on the JASPAR database^55^. Opposite regulation directions of the activation and silencing models were finally compared in terms of DEGs. All statistical analyses were performed by R programming. Raw RNA-seq data and gene count quantification were submitted to NCBI GEO data base with accession ID GSE155981 (https://www.ncbi.nlm.nih.gov/geo/query/acc.cgi?acc=GSE155981).

### Statistics

For analysis of serum biochemistry between two groups, a two-tailed t-test was performed. For analysis of cell counts, such as proliferating BECs, a Mann-Whitney U Test was performed. A p<0.05 was considered significant, and plots are mean ± SD. Detailed statistic information for each assay is in the figure legend. All statistical analysis and graph generation was performed using GraphPad Prism 7 software.

## Acknowledgements

This work was supported by NIH grants 1R01DK62277, 1R01DK100287, 1R01DK116993, R01CA204586, 1R01CA251155-01 and Endowed Chair for Experimental Pathology to S.P.M. This work was also supported in part by 1R01CA258449 to SK. This work was also supported by T32EB0010216 and 1F31DK115017-01 to J.O.R. This work was also supported by P30DK120531 to the Pittsburgh Liver Research Center for services provided by Biospecimen Repository and Processing Core.

## Contributor

Conceived project, analyzed and interpreted results, wrote manuscript: SH, JOR, SK, SPM

Performed experiments: SH, JOR, RR, KK, AB, SS, EH,

Provided reagents or performed specialized analysis and interpretations: SL, SS, MP, AB, DS, RR, ADS, KN, JT

---

**Supplementary Table 1:** Patient samples used in the study.

| **Identifier** | **Age** | **Sex** | **Diagnosis** | **Miscellaneous** |
| --- | --- | --- | --- | --- |
| PHS16-32066 | 66 | M | Non-neoplastic hepatic parenchyma | Mild nodular regenerative hyperplasia and mild portal reactive changes |
| PHS16-40535 | 50 | M | Non-neoplastic hepatic parenchyma | Minimal mixed micro-and macrovesicular steatosis involving approximately 5% of the hepatocytes. Mild chronic portal inflammation. Focal mild portal fibrosis. |
| PHS18-5904 | 71 | M | Background liver | Mild portal inflammation. No significant steatosis or fibrosis |
| PHS18-11592 | 60 | F | Background liver | Moderate macrovesicular steatosis |
| PHS18-34295 | 53 | M | Non-neoplastic hepatic parenchyma | Mild nodular regenerative hyperplasia |
| PHS18-4910 | 48 | M | Alcoholic hepatitis | Micronodular liver cirrhosis with moderate cholestasis, ductular reaction, abundant Mallory-Denk bodies, neutrophilic lobular inflammation, compatible with clinical history of acute alcoholic hepatitis. Chronic active steatohepatitis (Nash score: 5/8), 15% mixed micro-and macrovesicular steatosis. |
| PHS18-12144 | 72 | M | Alcoholic hepatitis | Active mixed cirrhosis secondary to chronic hepatitis, viral-type C, mildly active. Marked cholestasis and focal parenchymal extinction with prominent ductular reaction. |
| PHS18-42469 | 41 | M | Alcoholic hepatitis | micronodular cirrhosis with extensive parenchymal extinction and ductular reaction, secondary to alcohol abuse. mixed micro-macrovesicular steatosis involving about 50% of hepatocytes. |
| PHS19-24672 | 61 | M | Alcoholic hepatitis | mixed micro- and macronodular cirrhosis. clinical history of ethanol use. areas of parenchymal extinction with marked ductular reaction replacement and focal cholestasis. |
| PHS19-31767 | 63 | M | Nonalcoholic steatohepatitis (NASH) | Mixed macro and micronodular cirrhosis clinically secondary to NASH (Nash activity score: 2/8) (fibrosis stage: 4/4). Areas of parenchymal extinction with marked ductular reaction replacement and focal cholestasis. |
| PHS19-32169 | 66 | M | Alcoholic hepatitis | Macronodular cirrhosis, clinically due to Hepatitis C and alcoholic steatohepatitis. Focal areas of parenchymal extinction with marked ductular reaction replacement. |
| PHS19-40351 | 42 | M | Alcoholic hepatitis | Predominantly micronodular cirrhosis secondary to chronic steatohepatitis occurring in setting of obesity and ethanol use. Areas of parenchymal extinction with marked ductular reaction. Hepatocyte ballooning with Mallory-Denk bodies, neutrophils and hepatocanalicular cholestasis. Macrovesicular steatosis involving approximately 30-40% of hepatocytes. |
| PHS20-9506 | 54 | F | NASH | Mixed micro- and macronodular cirrhosis; clinical non-alcoholic steatohepatitis. a. numerous alpha-1 antitrypsin globules in periportal hepatocytes highlighted by pas/d and AAT immunostain. Mild mixed steatosis involving approximately 5% of hepatocytes. Areas of parenchymal extinction with marked ductular reaction replacement. |
| PHS20-9600 | 56 | F | Alcoholic hepatitis | Mixed micro- and macronodular cirrhosis with minimal residual steatosis involving <5% of hepatocytes. Clinical history of non-alcoholic steatohepatitis. Areas of parenchymal extinction with marked ductular reaction replacement and focal cholestasis. |
| PHS20-10786 | 39 | F | Alcoholic hepatitis | Predominantly micronodular cirrhosis with large areas of parenchymal extension and florid ductular reaction. Clinically decompensated cirrhosis due to ethanol use. Focal ballooning degeneration with rare, poorly-formed, Mallory-Denk bodies and mega-mitochondria. |
| PHS16-44914 | 64 | F | Alcoholic hepatitis | Active mixed cirrhosis with focal parenchymal extinction. Focally severe mixed micro-macrovesicular steatosis involving 60-70% of hepatocytes with superimposed steatohepatitis, easily identifiable Mallory-Denk bodies and cholangiolar cholestasis. Consistent with clinical history of alcohol use. |
| PHS17-14821 | 62 | M | Alcoholic hepatitis | Active mixed but predominantly micronodular cirrhosis with residual mixed steatosis and occasional Mallory's hyaline deposition. Clinical history of alcohol-use. Occasional regenerative nodules with brisk ductular reaction. |
| PHS16-28155 | 67 | F | Polycystic liver disease | Benign cysts lined by biliary epithelium. Surrounding liver parenchyma with chronic inflammation, patchy scarring and multiple biliary hamartomas (Von Meyenberg complexes). Mild macrovesicular steatosis involving about 10% of hepatocytes. Findings consistent with clinical history of polycystic liver and kidney disease. |
| PHS12-30089 | 60 | F | Polycystic liver disease | Multiple biliary type cysts, consistent with the clinical history of polycystic liver disease. Vascular congestion and mild chronic inflammation of the cyst wall. Small islands of hepatocytes with sinusoidal congestion and mild microvesicular steatosis |
| PHS15-21076 | 35 | F | Polycystic liver disease | Polycystic liver disease with Von Meyenberg complexes. Nodular regenerative hyperplasia and mild non-specific portal as well as lobular inflammation. |
| PHS12-35452 | 50 | F | Polycystic liver disease | Multiple biliary cysts, consistent with polycystic liver disease. Extensive hemorrhage, chronic inflammation, and fibrosis in the wall of biliary cysts. |
| PHS10-4073 | 48 | F | Polycystic liver disease | Multiple biliary cysts and multiple Von Meyenburg's complexes, consistent with the clinical history of adult polycystic kidney and liver disease. |
| PHS12-34148 | 34 | F | Cystic fibrosis | Bile ductular proliferation with cholangiolitis. Portal fibrosis with focal portal to portal early fibrous bridge formation. Histologic changes are nonspecific and are compatible with the underlying diagnosis of cystic fibrosis. |
| PHS15-14513 | 32 | M | Cystic fibrosis | Large areas of fibrosis with thick fibrous bands suggestive of marked architectural distortion. Numerous occasional hepatocytes with pseudo-ground glass cytoplasm. Architectural distortion with fibrous bands. The clinical history of lung transplantation secondary to cystic fibrosis is noted. The pattern of fibrosis development in patients with cystic fibrosis can be irregular and focal, leading to incomplete hepatic fibrosis (focal biliary cirrhosis/fibrosis). |
| PHS16-1251 | 23 | F | Cystic fibrosis | Intact hepatic architecture with diffusely increased hepatocyte glycogen. Mild diminution in portal vein caliber. Mild microvesicular steatosis. Mild increase in reticuloendothelial iron stores History of elevated alkaline phosphatase, AST, GGTP and ammonia with normal bilirubin and ALT levels who underwent double lung transplant in november 2015 for treatment of cystic fibrosis. |
| PHS17-19070 | 23 | M | Cystic fibrosis | Prominent macrovesicular steatosis involving approximately 90% of sampled liver parenchyma with superimposed minimally active steatohepatitis (NAS active score = 4-5/8). Portal, periportal and pericellular fibrosis with scattered delicate non-bridging fibrous septae (fibrosis stage = 2-3/4). Focal marked ductular proliferation with associated cholangiolitis, ductular ectasia and inspissated luminal secretions. Findings consistent with the clinical history of cystic fibrosis |
| PHS17-35744 | 22 | M | Cystic fibrosis | Cystic fibrosis related liver disease with PAS-positive inspissated bile plugs and patchy periportal/sinusoidal fibrosis (fibrosis stage 2/4). Bile duct dilatation with cholangiolar proliferation, neutrophilic cholangitis, lobular ballooning degeneration and canalicular cholestasis. Consistent with cystic fibrosis. |
| TP10-P531 | 31 | F | Cystic fibrosis | Decompensated cirrhosis in the setting of cystic fibrosis. Cholangiolitis with marked ductular proliferation, ductular ectasia and biliary sludge consistent with CF. Severe mixed micro- and macro-vesicular steatosis. NAS=6/8 and fibrosis stage=4/4 |

**Supplementary Table 2:**
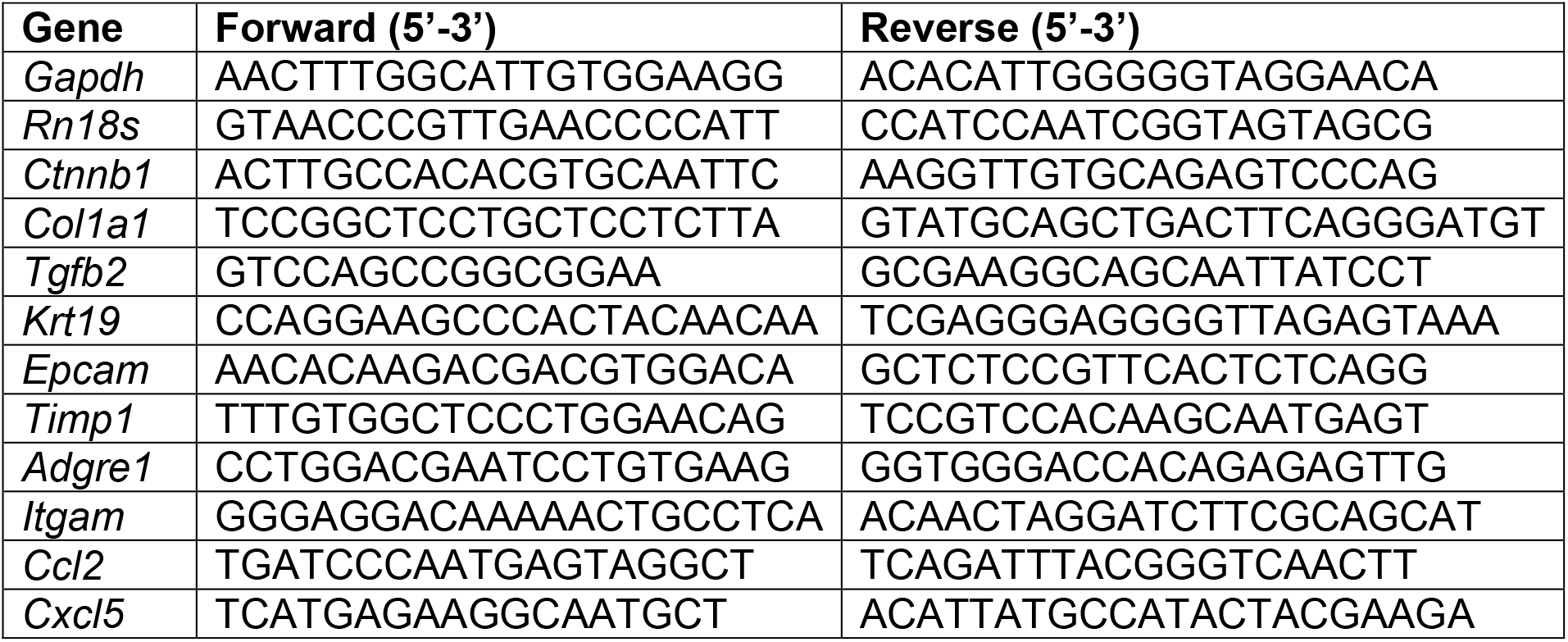
Sequence of qPCR primers.

## SUPPLEMENTARY FIGURES AND FIGURE LEGENDS

**Figure S1:**
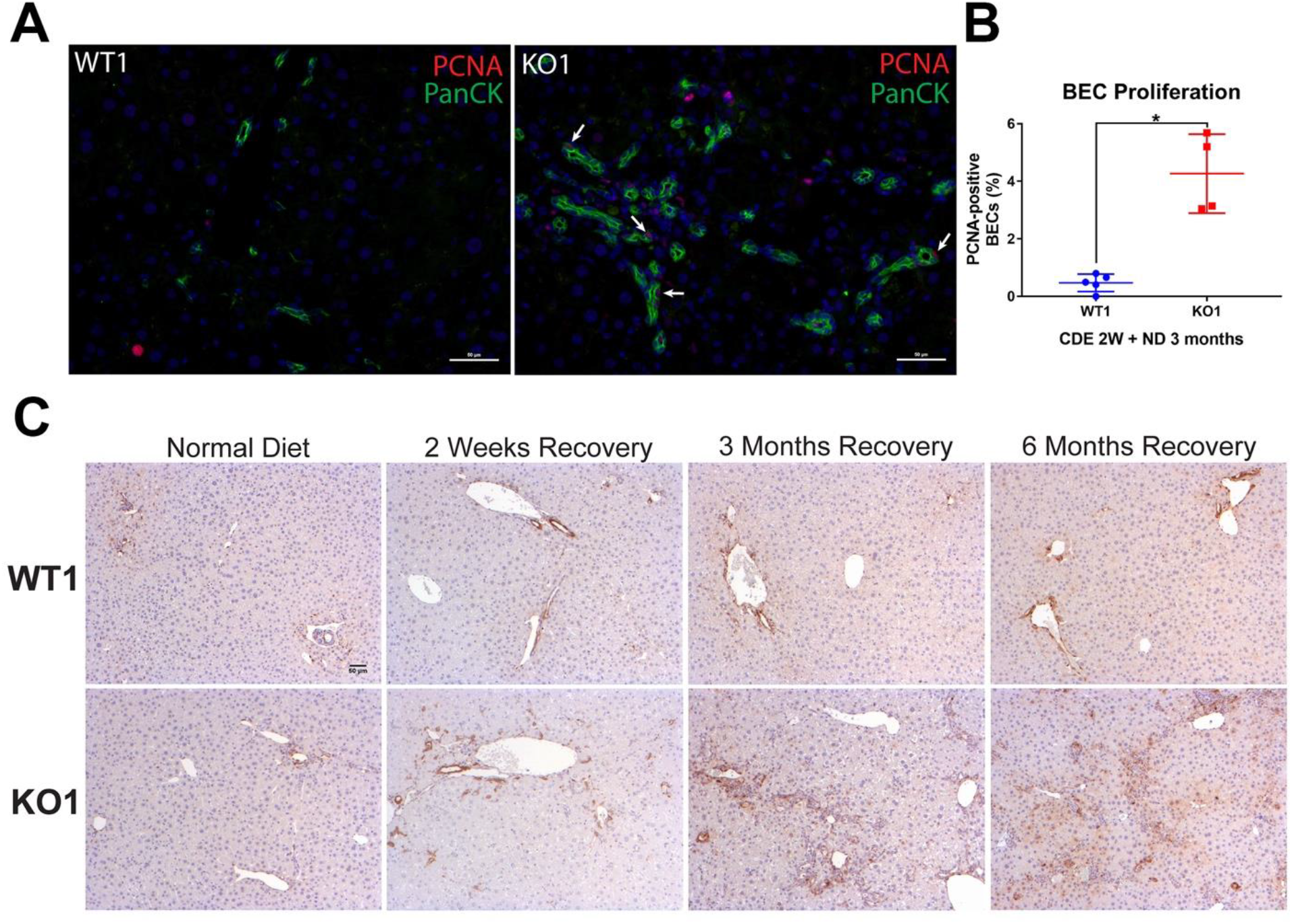
Enhanced PCNA and increased p-Erk staining in BECs in KO1 during recovery on normal diet after initial 2w CDE diet induced injury. A) PanCK (green) and PCNA (red) immunofluorescence in WT1 and KO1 mice at 3m of recovery on normal diet after 2w CDE diet. Scale bar = 50µm. B) Quantification of PCNA-positive BECs at 3m recovery on normal diet (ND) from 2w CDE (one-way ANOVA, *p<0.05). C) WT1 and KO1 mice staining with an antibody against phospho-Erk1/2 (Thr202/Tyr204) reveals staining in a subset of BECs in the ductular reaction in KO1 mice. Scale bar = 50µm.

**Figure S2.**
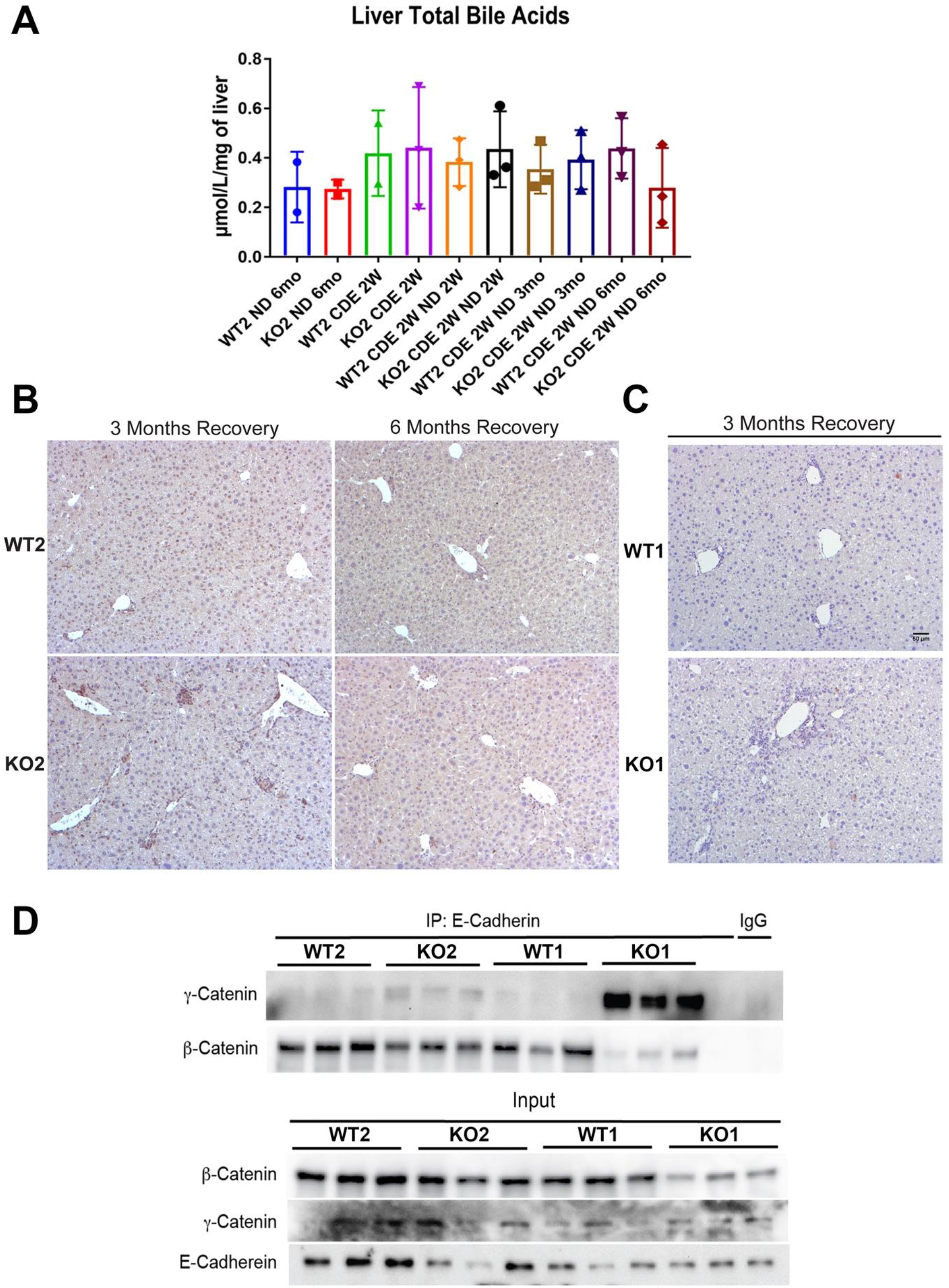
Bile acids and cell death are not the basis of fibrosis and ductular reaction due to CDE diet, and differences in adherens junction integrity don’t explain phenotypic differences between KO2 and KO1 at 6m of recovery. A) Quantification of bile acids in whole livers in WT2 and KO2 at 2w after CDE diet and at various time recovery times on normal diet. B) p21 staining shows no positive cells at 3m or 6m of recovery on normal diet in WT2 and KO2 liver sections. C) Cleaved caspase 3 staining reveals almost no ongoing cell death in the livers of WT1 or KO1 mice after 3m of recovery from CDE diet-induced liver injury. Scale bar = 50µm. D) Immunoprecipitation studies show E-cadherin association with β-catenin in WT1, WT2 and KO2 livers at 6m of recovery while it associates with γ-catenin in KO1 at the same time due to continued lack of β-catenin in KO1 (top panels). Input verifies low levels of β-catenin in whole liver lysates of KO1 at the same time depicting β-catenin presence in liver non-epithelial cells (bottom panels).

**Figure S3.**
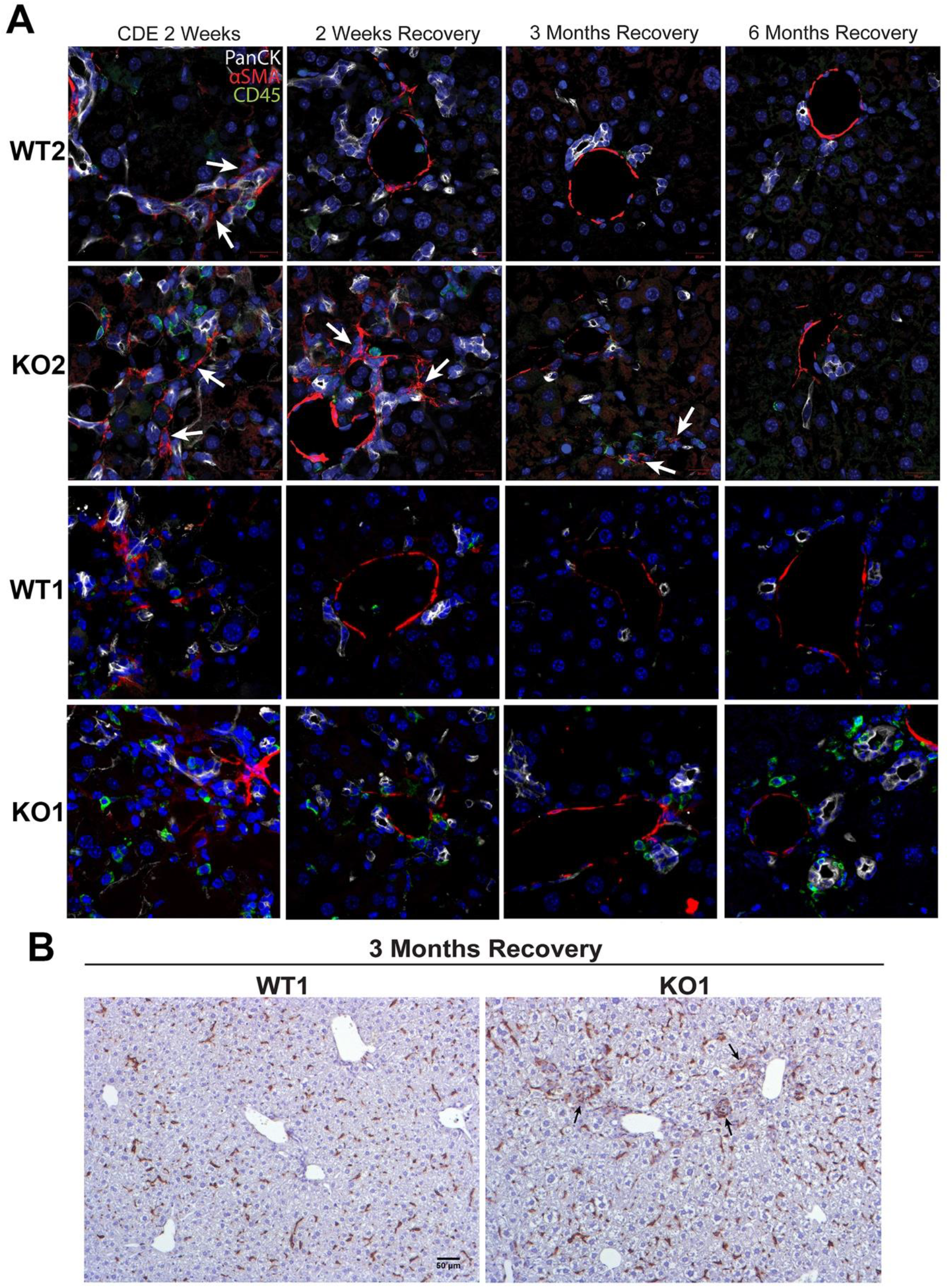
Immune cells continue to prevail in periportal region in KO1 even at 6m of recovery while they subside in all other genotypes after 3m of recovery or earlier. A) Representative confocal image of triple immunofluorescence for PanCK (white), αSMA (red), and CD45 (green) in WT1, KO1, WT2 and KO2 at 2w of CDE diet and recovery on normal diet for 2w, 3m or 6m. Cells expressing αSMA are closely associated with PanCK-positive cells (white arrows). Scale bar = 20µm. B) F4/80 staining reveals macrophages closely associated with the DR in KO1 mice at 3m of recovery (black arrows). Scale bar = 50µm.

**Figure S4:**
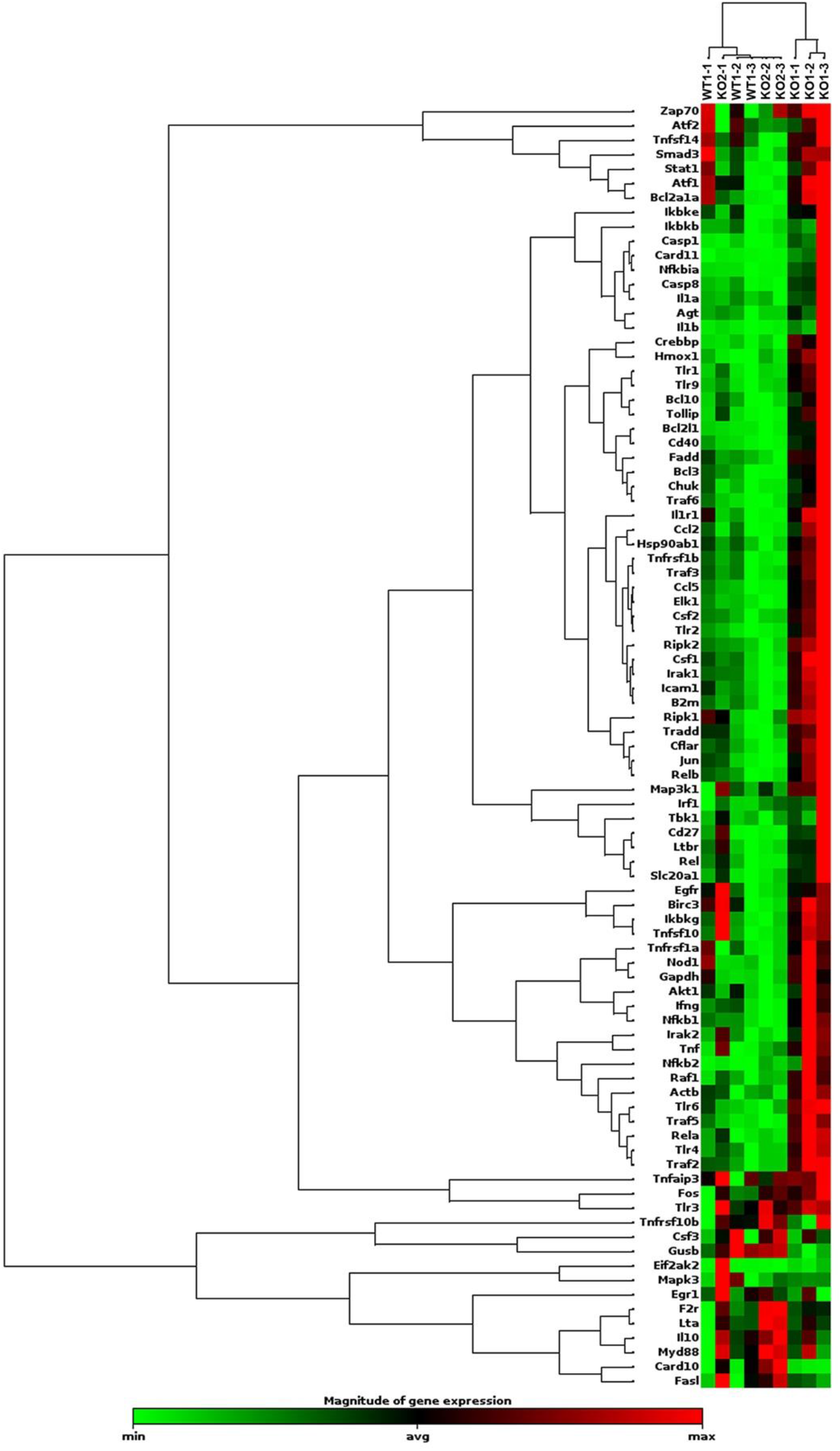
Evidence of NF-κB activation in KO1 livers but not in WT1 or KO2 livers at 6m of recovery from CDE diet. RNA from whole livers of WT1, KO1 and KO2 (n=3 each) at 6m recovery was assessed for 84 NF-κB downstream target genes by RT-PCR array. Using a fold change threshold = 2, p-value threshold = 0.05, we identified several genes altered in KO1 only and clustergram showed clear separation of KO1 from KO2 and WT1.

**Figure S5:**
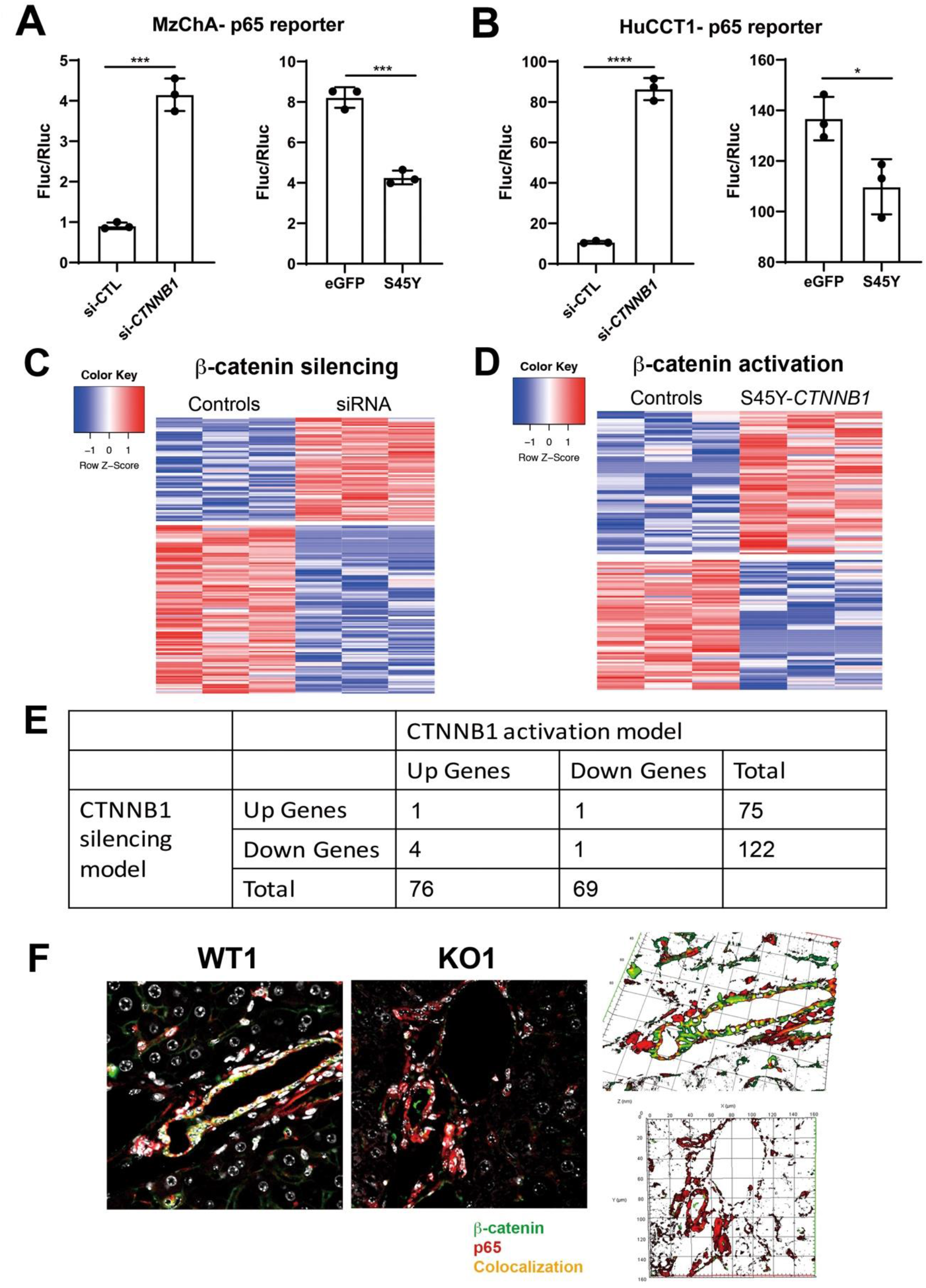
Modulation of β-catenin in cholangiocytes impacts NF-κB activity due to p65-β-catenin complex. A) Reporter assay shows knockdown of *Ctnnb1* stimulates p65 transcriptional activity (left) while transfection of stable S45Y-β-catenin represses p65 reporter activity (right) in MzChA human cholangiocarcinoma cells (Unpaired t-test, ***p<0.001). B) Reporter assay shows knockdown of *Ctnnb1* stimulates p65 transcriptional activity (left) while expression of constitutively active S45Y-β-catenin suppresses p65 transcriptional activity (right) in HuCCT1 human cholangiocarcinoma cells (Unpaired t-test, *p<0.05, **p<0.01, ****p<0.0001). C) Heatmap of the differentially expressed genes in *Ctnnb1* loss-of-function in SMCC. D) Heatmap of the differentially expressed genes in *CTNNB1* gain-of-function in SMCC. E) Very few common differentially expressed genes between *Ctnnb1* loss- and gain-of-function models were identified, although altogether 335 genes were altered. These genes were assessed for transcription factor (TF) binding profiles by JASPAR (presented in Fig.7E). F) Representative confocal images showing the expression of p65 (red), β-catenin (green) and DAPI (white) in WT1 and KO1 livers (left). To visualize colocalization for p65 (red) and β-catenin (green), 3D images were reconstructed using Zen blue 2012 software (right). The reconstruction was performed with 9 confocal z-stacks with 0.8 µm increments and yellow region indicates colocalization of p65 (red) and β-Catenin (green) in the projection on the X, Y and Z planes.

## REFERENCES

1. Pellicoro A, Ramachandran P, Iredale JP, Fallowfield JA. Liver fibrosis and repair: immune regulation of wound healing in a solid organ. Nat Rev Immunol 14, 181–194 (2014).

2. Asrani SK, Devarbhavi H, Eaton J, Kamath PS. Burden of liver diseases in the world. J Hepatol 70, 151–171 (2019).

3. Argemi J, et al. Defective HNF4alpha-dependent gene expression as a driver of hepatocellular failure in alcoholic hepatitis. Nat Commun 10, 3126 (2019).

4. Nishikawa T, et al. Resetting the transcription factor network reverses terminal chronic hepatic failure. J Clin Invest 125, 1533–1544 (2015).

5. Sato K, Marzioni M, Meng F, Francis H, Glaser S, Alpini G. Ductular Reaction in Liver Diseases: Pathological Mechanisms and Translational Significances. Hepatology 69, 420–430 (2019).

6. Wilson DB, Rudnick DA. Invasive Ductular Reaction: Form and Function. Am J Pathol 189, 1501–1504 (2019).

7. Lowes KN, Brennan BA, Yeoh GC, Olynyk JK. Oval cell numbers in human chronic liver diseases are directly related to disease severity. Am J Pathol 154, 537–541 (1999).

8. Richardson MM, et al. Progressive fibrosis in nonalcoholic steatohepatitis: association with altered regeneration and a ductular reaction. Gastroenterology 133, 80–90 (2007).

9. Zhao L, Westerhoff M, Pai RK, Choi WT, Gao ZH, Hart J. Centrilobular ductular reaction correlates with fibrosis stage and fibrosis progression in non-alcoholic steatohepatitis. Mod Pathol 31, 150–159 (2018).

10. Planas-Paz L, et al. YAP, but Not RSPO-LGR4/5, Signaling in Biliary Epithelial Cells Promotes a Ductular Reaction in Response to Liver Injury. Cell Stem Cell 25, 39–53.e10 (2019).

11. Apte U, et al. Beta-catenin activation promotes liver regeneration after acetaminophen-induced injury. Am J Pathol 175, 1056–1065 (2009).

12. Tan X, Behari J, Cieply B, Michalopoulos GK, Monga SP. Conditional deletion of beta-catenin reveals its role in liver growth and regeneration. Gastroenterology 131, 1561–1572 (2006).

13. Akhurst B, et al. A modified choline-deficient, ethionine-supplemented diet protocol effectively induces oval cells in mouse liver. Hepatology 34, 519–522 (2001).

14. Russell JO, et al. Hepatocyte-Specific β-Catenin Deletion During Severe Liver Injury Provokes Cholangiocytes to Differentiate Into Hepatocytes. Hepatology 69, 742–759 (2019).

15. Yoshiji H, et al. Tissue inhibitor of metalloproteinases-1 promotes liver fibrosis development in a transgenic mouse model. Hepatology 32, 1248–1254 (2000).

16. Yoshiji H, et al. Tissue inhibitor of metalloproteinases-1 attenuates spontaneous liver fibrosis resolution in the transgenic mouse. Hepatology 36, 850–860 (2002).

17. Pepe-Mooney BJ, et al. Single-Cell Analysis of the Liver Epithelium Reveals Dynamic Heterogeneity and an Essential Role for YAP in Homeostasis and Regeneration. Cell Stem Cell 25, 23–38.e28 (2019).

18. Fickert P, Wagner M. Biliary bile acids in hepatobiliary injury - What is the link? J Hepatol 67, 619–631 (2017).

19. Behari J, et al. Liver-specific beta-catenin knockout mice exhibit defective bile acid and cholesterol homeostasis and increased susceptibility to diet-induced steatohepatitis. Am J Pathol 176, 744–753 (2010).

20. Thompson MD, et al. beta-Catenin regulation of farnesoid X receptor signaling and bile acid metabolism during murine cholestasis. Hepatology 67, 955–971 (2018).

21. Lu WY, et al. Hepatic progenitor cells of biliary origin with liver repopulation capacity. Nat Cell Biol 17, 971–983 (2015).

22. Pradhan-Sundd T, et al. Dysregulated Bile Transporters and Impaired Tight Junctions During Chronic Liver Injury in Mice. Gastroenterology 155, 1218–1232 e1224 (2018).

23. Pradhan-Sundd T, et al. Dual catenin loss in murine liver causes tight junctional deregulation and progressive intrahepatic cholestasis. Hepatology 67, 2320–2337 (2018).

24. Knight B, et al. Attenuated liver progenitor (oval) cell and fibrogenic responses to the choline deficient, ethionine supplemented diet in the BALB/c inbred strain of mice. J Hepatol 46, 134–141 (2007).

25. Guillot A, Tacke F. Liver Macrophages: Old Dogmas and New Insights. Hepatol Commun 3, 730–743 (2019).

26. Nejak-Bowen K, Kikuchi A, Monga SP. Beta-catenin-NF-kappaB interactions in murine hepatocytes: a complex to die for. Hepatology 57, 763–774 (2013).

27. Fava G, Glaser S, Francis H, Alpini G. The immunophysiology of biliary epithelium. Semin Liver Dis 25, 251–264 (2005).

28. Pinto C, Giordano DM, Maroni L, Marzioni M. Role of inflammation and proinflammatory cytokines in cholangiocyte pathophysiology. Biochim Biophys Acta Mol Basis Dis 1864, 1270–1278 (2018).

29. Kim KH, Chen CC, Alpini G, Lau LF. CCN1 induces hepatic ductular reaction through integrin alphavbeta(5)-mediated activation of NF-kappaB. J Clin Invest 125, 1886–1900 (2015).

30. Luedde T, Schwabe RF. NF-kappaB in the liver--linking injury, fibrosis and hepatocellular carcinoma. Nat Rev Gastroenterol Hepatol 8, 108–118 (2011).

31. Chang XB, et al. Role of N-linked oligosaccharides in the biosynthetic processing of the cystic fibrosis membrane conductance regulator. J Cell Sci 121, 2814–2823 (2008).

32. Nejak-Bowen K. If It Looks Like a Duct and Acts Like a Duct: On the Role of Reprogrammed Hepatocytes in Cholangiopathies. Gene Expr 20, 19–23 (2020).

33. Kamimoto K, Nakano Y, Kaneko K, Miyajima A, Itoh T. Multidimensional imaging of liver injury repair in mice reveals fundamental role of the ductular reaction. Commun Biol 3, 289 (2020).

34. Raven A, et al. Cholangiocytes act as facultative liver stem cells during impaired hepatocyte regeneration. Nature 547, 350–354 (2017).

35. Aguilar-Bravo B, et al. Ductular Reaction Cells Display an Inflammatory Profile and Recruit Neutrophils in Alcoholic Hepatitis. Hepatology 69, 2180–2195 (2019).

36. Deng J, et al. beta-catenin interacts with and inhibits NF-kappa B in human colon and breast cancer. Cancer Cell 2, 323–334 (2002).

37. Russell JO, Monga SS. Wnt/β-Catenin Signaling in Liver Development, Homeostasis, and Pathobiology. Annu Rev Pathol, (2017).

38. Apte U, Thompson MD, Cui S, Liu B, Cieply B, Monga SP. Wnt/beta-catenin signaling mediates oval cell response in rodents. Hepatology 47, 288–295 (2008).

39. Hu M, et al. Wnt/beta-catenin signaling in murine hepatic transit amplifying progenitor cells. Gastroenterology 133, 1579–1591 (2007).

40. Okabe H, et al. Wnt signaling regulates hepatobiliary repair following cholestatic liver injury in mice. Hepatology 64, 1652–1666 (2016).

41. O’Hara SP, Tabibian JH, Splinter PL, LaRusso NF. The dynamic biliary epithelia: molecules, pathways, and disease. J Hepatol 58, 575–582 (2013).

42. Wilson DH, et al. Non-canonical Wnt signalling regulates scarring in biliary disease via the planar cell polarity receptors. Nat Commun 11, 445 (2020).

43. Sakiani S, Kleiner DE, Heller T, Koh C. Hepatic Manifestations of Cystic Fibrosis. Clin Liver Dis 23, 263–277 (2019).

44. Cohn JA, Strong TV, Picciotto MR, Nairn AC, Collins FS, Fitz JG. Localization of the cystic fibrosis transmembrane conductance regulator in human bile duct epithelial cells. Gastroenterology 105, 1857–1864 (1993).

45. Pankow S, et al. F508 CFTR interactome remodelling promotes rescue of cystic fibrosis. Nature 528, 510–516 (2015).

46. Liu K, Zhang X, Zhang JT, Tsang LL, Jiang X, Chan HC. Defective CFTR- beta-catenin interaction promotes NF-kappaB nuclear translocation and intestinal inflammation in cystic fibrosis. Oncotarget 7, 64030–64042 (2016).

47. Fiorotto R, et al. The cystic fibrosis transmembrane conductance regulator controls biliary epithelial inflammation and permeability by regulating Src tyrosine kinase activity. Hepatology 64, 2118–2134 (2016).

48. Ko S, Russell JO, Molina LM, Monga SP. Liver Progenitors and Adult Cell Plasticity in Hepatic Injury and Repair: Knowns and Unknowns. Annu Rev Pathol, (2019).

49. Leroy V, et al. Circulating matrix metalloproteinases 1, 2, 9 and their inhibitors TIMP-1 and TIMP-2 as serum markers of liver fibrosis in patients with chronic hepatitis C: comparison with PIIINP and hyaluronic acid. Am J Gastroenterol 99, 271–279 (2004).

50. Boeker KH, Haberkorn CI, Michels D, Flemming P, Manns MP, Lichtinghagen R. Diagnostic potential of circulating TIMP-1 and MMP-2 as markers of liver fibrosis in patients with chronic hepatitis C. Clin Chim Acta 316, 71–81 (2002).

51. Bolger AM, Lohse M, Usadel B. Trimmomatic: a flexible trimmer for Illumina sequence data. Bioinformatics 30, 2114–2120 (2014).

52. Kim D, Langmead B, Salzberg SL. HISAT: a fast spliced aligner with low memory requirements. Nat Methods 12, 357–360 (2015).

53. Anders S, Pyl PT, Huber W. HTSeq--a Python framework to work with high-throughput sequencing data. Bioinformatics 31, 166–169 (2015).

54. Love MI, Huber W, Anders S. Moderated estimation of fold change and dispersion for RNA-seq data with DESeq2. Genome Biol 15, 550 (2014).

55. Fornes O, et al. JASPAR 2020: update of the open-access database of transcription factor binding profiles. Nucleic Acids Res 48, D87–D92 (2020).

